# YAP activation reverses aging-related visual dysfunction caused by impaired cell‒matrix adhesion

**DOI:** 10.64898/2026.01.07.697636

**Authors:** Gyuri Kim, Chanok Son, Hyo Kyung Lee, Jaemyung Choi, Soomin Lee, Seyoun Oh, Jae-Byoung Chae, Chul-Woo Park, Chaekyu Kim, Ja-Hyoung Ryu, Semin Lee, Jiwon Jang, Hyewon Chung

**Author notes:** Contributed equally (Gyuri Kim, Chanok Son, Hyo Kyung Lee). Correspondence: Semin Lee, Jiwon Jang, Hyewon Chung.

## Abstract

Cellular senescence of retinal pigment epithelium (RPE) cells drives age-related visual decline, particularly in the pathology of age-related macular degeneration (AMD). While genome-wide association studies (GWAS) have identified genetic risk factors underlying AMD, the molecular mechanisms governing RPE senescence remain unclear. Here, single-cell RNA sequencing of young and old mouse RPE revealed dysregulated cell-matrix adhesion as a key feature of senescence, consistent with transcriptional changes in AMD patients. Hydrogel-based experiments confirmed that impaired integrin-mediated adhesion induces RPE senescence. Yes-associated protein 1 (YAP), a crucial mechanotransducer, mediated the protective effects of cell-matrix adhesion, and its activation alone reversed aging phenotypes in senescent RPE cells. Notably, treatment with TRULI, a small-molecule YAP activator, significantly improved visual function in AMD and naturally aged mice. These findings highlight the integrin-YAP mechanotransduction pathway as a fundamental regulator of RPE senescence and a potential therapeutic target for AMD.

## INTRODUCTION

The retinal pigment epithelium (RPE), an epithelial cell monolayer positioned between the photoreceptors and choriocapillaris, is crucial for maintaining retinal health and visual function^1,2^. RPE dysfunction is closely associated with the onset and progression of age-related macular degeneration (AMD), the leading cause of vision loss globally^3,4^. Although anti-vascular endothelial growth factor (anti-VEGF) treatments effectively manage wet AMD (neovascular), they require repeated intravitreal injections and do not fully halt disease progression. Until recently, no effective treatments were available for the more prevalent form of AMD, known as dry AMD (atrophic). Genome-wide association studies (GWAS) have identified single nucleotide polymorphisms (SNPs) associated with AMD, particularly in the complement system^5^. However, despite extensive research, complement-regulating agents targeting geographic atrophy (GA), the late stage of dry AMD, have not yielded successful outcomes. Although two complement inhibitors—a C3 inhibitor and a C5 inhibitor—have recently received FDA approval, they have not markedly improved vision^6,7^. The multifactorial nature of AMD suggests that targeting the complement pathway alone is insufficient, and this symptomatic approach fails to halt or reverse disease progression. These results highlight the need for safe and effective treatments that can be introduced earlier in the disease process and that target the underlying causes of AMD, such as cellular aging.

AMD development primarily results from age-related RPE deterioration^8,9^. Cellular senescence, a state of irreversible cell cycle arrest triggered by various stresses, leads to characteristic changes, such as enlarged cell size, accumulation of senescence-associated β-galactosidase (SA-β-Gal), and increased proinflammatory cytokine, chemokine, growth factor, and protease secretion, collectively termed the senescence-associated secretory phenotype (SASP)^10–12^. Senescent RPE cells contribute to retinal degeneration, particularly under conditions such as AMD, and targeting these cells with senolytic drugs can mitigate this degeneration^13–15^. However, the precise mechanisms underlying RPE cell senescence remain unclear. Understanding these mechanisms is crucial for advancing AMD research and developing innovative treatments.

To investigate the mechanisms underlying RPE senescence, we performed single-cell RNA sequencing (scRNA-seq) on RPE cells from young and old mice. Although previous studies have illuminated the single-cell transcriptomic changes in various retinal cells with age^16,17^, these studies did not specifically focus on the RPE, largely because of the technical challenges associated with isolating these cells and their relatively low abundance. Our scRNA-seq analysis uncovered a significant impairment in cell-matrix adhesion in aged RPE cells, closely aligning with AMD pathology, characterized by the accumulation of drusen—extracellular deposits between the RPE and Bruch’s membrane (BrM). Drusen, composed of lipids, apolipoproteins, complement factors, and RPE-derived cellular debris, plays a central role in AMD progression by triggering inflammatory responses, activating complement pathways, and disrupting nutrient and waste exchange between the choroid and retina^3,18,19^. Notably, drusen can directly interfere with the physical interaction between the RPE and BrM. However, the functional impact of impaired cell-matrix interaction on RPE senescence remains unknown.

In this study, we investigated the role of cell-matrix adhesion in RPE senescence using hydrogel systems and genetic and chemical tools. We found that dysfunctions in integrin-mediated adhesion are a key driver of RPE senescence. Mechanistically, our work identifies Yes-associated protein 1(YAP), a stem cell factor known for its ability to reprogram differentiated cells into tissue-specific stem cells^20^, as a critical link between cell-matrix adhesion and the onset of RPE senescence. Similar to reprogramming factors like OCT4, SOX2, KLF4, and MYC (OSKM) that ameliorate age-related phenotypes^21–25^, transient activation of YAP effectively reactivated the stem cell transcriptional program, reversing the senescent state of RPE cells. Moreover, TRULI^26^, a small-molecule YAP activator, significantly restored visual function in both an AMD mouse model and naturally aged mice. This research highlights the dysregulation of the integrin-YAP mechanotransduction pathway as a fundamental mechanism driving RPE senescence. Importantly, compared to the complexity of using a combination of multiple reprogramming factors, our findings suggest that YAP activation alone holds significant therapeutic potential, offering a promising strategy for reversing RPE aging and restoring visual function.

## RESULTS

### Impaired cell-matrix adhesion in the RPE of aged mice and AMD patients revealed by single-cell RNA sequencing

scRNA-seq was performed on RPE cells isolated from 3- and 24-month-old mice to identify transcriptomic changes in aged RPE cells (Figure 1A). In total, 27,840 cells were categorized into 20 distinct clusters using uniform manifold approximation and projection (UMAP) (Figure 1B). Each cluster was annotated based on representative marker genes for each cell type^27,28^. Clusters 2, 3, 6, 12, 14, and 15 were grouped as RPE cells with high *Rpe65* and *Ttr* expression (Figures 1B and S1A). Differentially expressed gene (DEG) analysis identified 21 upregulated (log_2_FC ≥ 0.5, adjP < 0.05) and 66 downregulated DEGs (log_2_FC ≤ -0.5, adjP < 0.05) in the RPE cells derived from old mice compared to those from young mice (Figure S1B and Table S1). Despite the relatively small number of DEGs, Gene Ontology (GO) analysis with downregulated DEGs revealed significant enrichment of GO terms for cellular components associated with cell-extracellular matrix (ECM) adhesion (Figures S1C and S1D). These results suggest the dysfunction of cell-ECM interaction in old RPE cells.

**Fig. 1.**
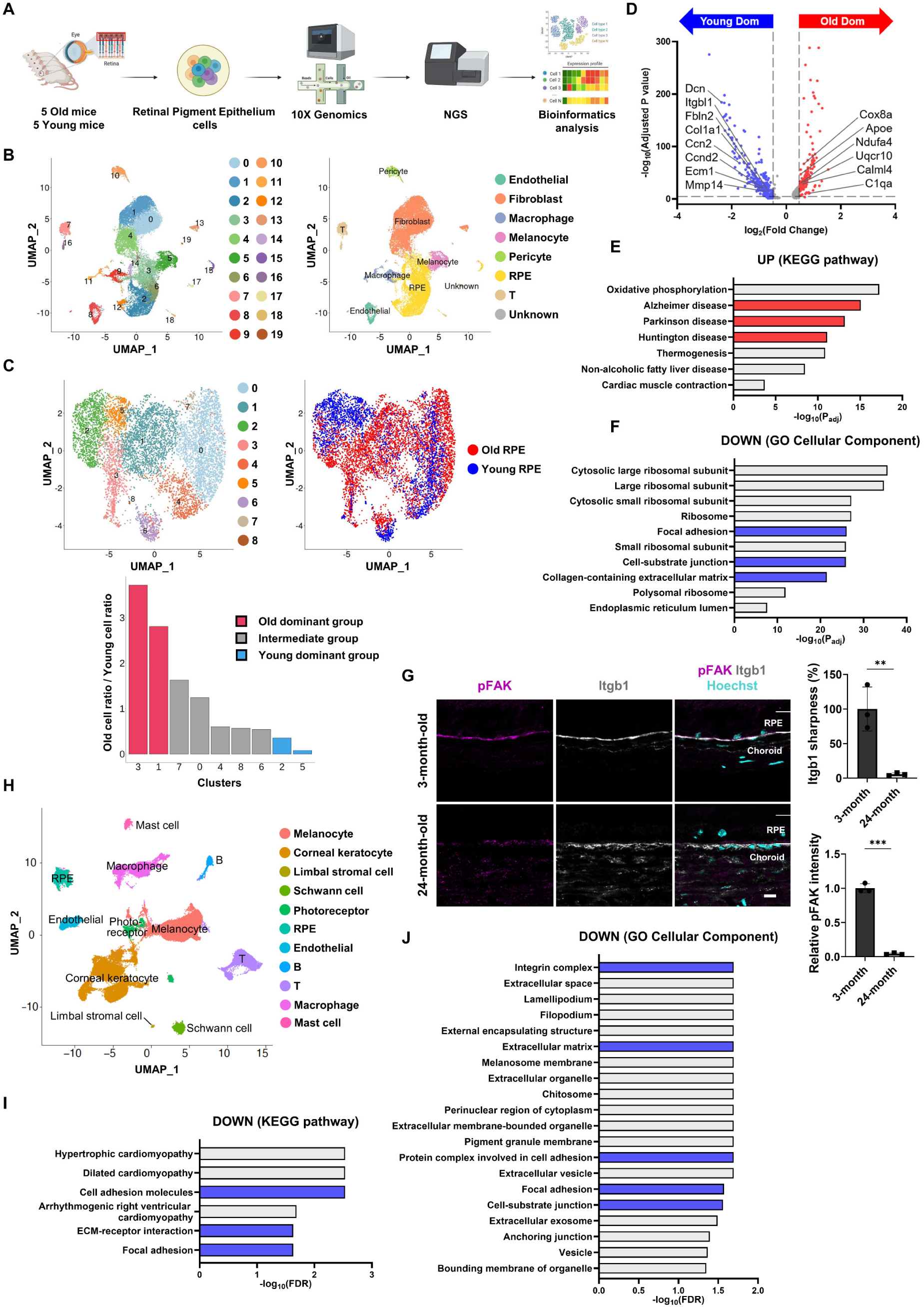
scRNA-seq reveals impaired cell-matrix adhesion in aged mouse and AMD patient RPE. (A) Schematic illustration of the scRNA-seq process for RPE cells from 3-month-old and 24-month-old mice. (B) UMAP analysis and cluster annotation of 27,840 scRNA-seq profiles. (C) UMAP analysis and distribution graph of old and young RPE cells grouped from b. (D) Volcano plot of DEGs from old-dominant and young-dominant clusters. The red and blue dots indicate either upregulated or downregulated genes in the old-dominant clusters, with cutoff values for DEGs: |log_2_FC| ≥ 0.5, adjP < 0.05. (E and F) The top pathways ranked by adjusted p-values from GO analysis with upregulated (E) and downregulated (F) genes from (D). (G) Immunostaining for pFAK and Itgb1 in cryosectioned retina/RPE/choroid tissues. The sharpness of Itgb1 was measured by the ratio of signal intensity to the width. *n* = 3 samples per group. (H) UMAP analysis and cluster annotation of 83,740 scRNA-seq profiles from fetal, normal adult, and AMD human eyes (GSE203499 and GSE210543). (I and J) The top pathways ranked by FDR from GO analysis with downregulated genes in AMD RPE compared to normal RPE. Two-sided Student’s t-test, ***P < 0.001, **P < 0.01. Exact *P* values are listed in Table S10. Scale bar: 10 μm (G).

The RPE cells were then re-clustered into 9 UMAP clusters to gain deeper insights. Old and young RPE cells showed different distributions, with clusters 1 and 3 being dominant in old RPE cells and clusters 2 and 5 being predominant in young RPE cells (Figure 1C). DEG analysis identified 181 upregulated (log_2_FC ≥ 0.5, adjP < 0.05) and 264 downregulated genes (log_2_FC ≤ -0.5, adjP < 0.05) in the old-dominant clusters compared to the young-dominant clusters (Figure 1D and Table S2). GO analysis of the upregulated DEGs revealed significant enrichment of Kyoto Encyclopedia of Genes and Genomes (KEGG) pathway terms associated with degenerative diseases and aging, such as Alzheimer’s, Parkinson’s, and Huntington’s disease, indicating aged transcriptomic features of RPE cells in the old-dominant clusters (Figure 1E). Conversely, the downregulated DEGs in the old-dominant clusters exhibited GO terms associated with ribosomes (Figure 1F). Defects in ribosome biogenesis have recently been reported as features of senescent cells^29^. Significant enrichment of GO terms related to cell‒ECM adhesion, such as focal adhesion, cell-substrate junction, and collagen-containing ECM, was observed in the downregulated DEGs of the old-dominant clusters (Figure 1F), consistent with the results obtained from the analysis of the entire RPE cell population (Figure S1D). Given the key role of integrins in cell-ECM adhesion and focal adhesion formation^30^, the old-dominant clusters exhibited markedly reduced enrichment scores for genes associated with integrin binding and collagen-containing ECM (Figure S1E). To validate these transcriptomic findings, we examined the expression patterns of Integrin β1 (Itgb1) and phosphorylated focal adhesion kinase (pFAK, the active form of FAK, which is a reliable indicator of integrin activity^31^) in cryosections of young and old retina/RPE/BrM. Itgb1 and pFAK were prominently localized at the BrM in the young mouse RPE, indicating robust cell-ECM interactions. In contrast, old mouse RPE cells exhibited diffuse and scattered Itgb1 expression with decreased pFAK expression (Figure 1G). Western blot analysis confirmed a significant reduction in pFAK levels in old mouse RPE cells (Figure S1F), indicating impaired integrin-mediated cell-ECM adhesion.

To determine whether transcriptomic changes in the aged mouse RPE reflect those occurring in the human RPE during AMD progression, we integrated and analyzed two human scRNA-seq datasets (GSE203499 and GSE210543) comprising fetal, normal adult, and AMD eyes (Table S3)^32^. A total of 83,740 cells were clustered and annotated into 11 cell types based on marker gene expression and comparisons with pre-annotated Tabula Sapiens eye scRNA-seq data (Figure 1H)^33^. The RPE cluster, consisting of 4,721 cells, was validated by *RPE65* and *BEST1* expression (Figure S1G). Next, we performed DEG analysis on RPE cells, comparing adult vs. fetal RPE and AMD vs. normal adult RPE. In the adult vs. fetal RPE comparison, 596 DEGs (log_2_FC ≤ -1, adjP < 0.05) were downregulated in adult RPE cells (Table S4). GO analysis of these genes revealed a significant enrichment of terms related to cell‒ECM interaction (Figure S1H), suggesting an age-related decline in RPE adhesion. In the AMD vs. normal adult RPE comparison, 70 DEGs (log_2_FC ≤ -1, adjP < 0.05) were downregulated in AMD RPE cells (Table S5). GO analysis of these genes also showed significant enrichment of cellular components and KEGG pathways related to cell‒ECM adhesion (Figures 1I and 1J). Key cell adhesion genes, including *ITGB1*, *ITGB8*, *ITGAV*, and *COL3A1*, were downregulated in AMD RPE (Table S5). Gene set enrichment analysis (GSEA) demonstrated significant enrichment of genes associated with ECM organization, integrin binding, and focal adhesion among the downregulated genes in AMD RPE cells (Figure S1I). These results highlight impaired cell‒ECM interactions as a defining characteristic of aged RPE cells in mice and humans, with substantial implications for AMD progression.

### Impaired cell-ECM adhesion as a driver of RPE senescence

Next, we aimed to explore the causal relationships between impaired cell‒ECM interactions and the aging process of RPE. Healthy RPE cells form a tight interaction with BrM, which is characterized by a very stiff mechanical property (7∼19 MPa)^34^. The strength of cell‒ECM interaction can be modulated by substrate stiffness, with cells exhibiting weaker ECM adhesion on softer substrates (Figure 2A)^35,36^. Compared to plastic culture plates (∼1 GPa), ARPE19 cells, a human RPE cell line, showed reduced levels of focal adhesion and phosphorylated myosin light chain (pMLC) on soft substrates (28 and 1.5 kPa) (Figures S2A and S2B). pMLC serves as a marker of contractile actomyosin, representing a reliable indicator of cellular mechanical stress levels. Importantly, SA-β-gal and p21 staining indicated that culturing ARPE19 cells on soft substrates significantly increased the number of senescent cells (Figures 2B and 2C). Additionally, the 5-ethynyl-2′-deoxyuridine (EdU) incorporation assay confirmed cell cycle arrest of SA-β-gal^+^ senescent cells (Figure 2B). These findings suggest that reduced cell‒ECM interaction drives cellular senescence in RPE cells.

**Fig. 2.**
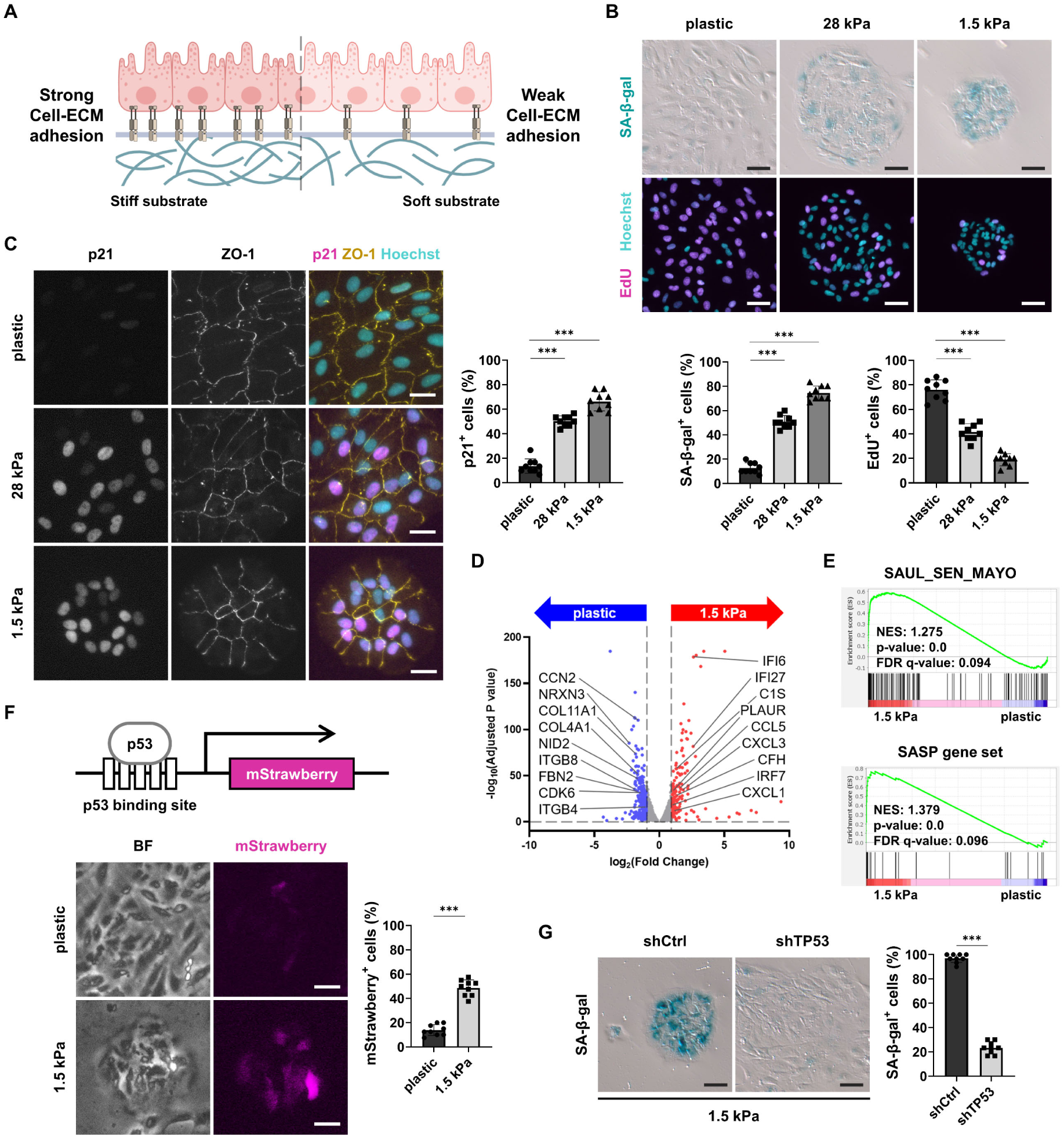
Impaired cell-ECM adhesion drives RPE senescence. (A) Schematic representation depicting the relationship between substrate stiffness and cell-ECM interaction. (B and C) SA-β-gal staining and EdU incorporation assay (B), and immunofluorescence assay for p21 (C) in ARPE19 cells cultured on substrate with different stiffness. *n* = 9 regions from three independent experiments. (D) Volcano plot of DEGs from ARPE19 cells cultured either on plastic or 1.5 kPa substrate. The red and blue dots indicate either upregulated or downregulated genes in ARPE19 cells on the 1.5 kPa substrate, with cutoff values for DEGs: |log_2_FC| ≥ 1, adjP < 0.05, average TPM value ≥ 5. (E) Gene set enrichment analysis (GSEA) with the RNA-seq data from ARPE19 cells cultured either on plastic or 1.5 kPa substrate. (F) ARPE19 cells expressing a p53 activity reporter cultured on plastic or 1.5 kPa for 5 days. *n* = 9 regions from three independent experiments. (G) SA-β-gal staining in ARPE19 cells cultured on 1.5 kPa hydrogel after TP53 KD. *n* = 9 regions from three independent experiments. Two-sided Student’s t-test, ***P < 0.001. Exact *P* values are listed in Table S10. Scale bars: 25 μm (C, F), 50 μm (B, G)

Bulk RNA-seq was performed on ARPE19 cells cultured on either plastic or 1.5 kPa substrates to investigate the transcriptome-wide effect of soft substrates (Figure 2D and Table S6). GO analysis with downregulated DEGs (log_2_FC ≤ -1, adjP < 0.05) at 1.5 kPa revealed enrichment of terms related to focal adhesion and ECM‒receptor interaction (Figure S2C), confirming the diminished cell‒ECM adhesion^37^. In contrast, upregulated DEGs at 1.5 kPa (log_2_FC ≥ 1, adjP < 0.05) included genes related to innate immune responses, with enrichment in interferon and cytokine signaling (Figure S2D). Moreover, GSEA demonstrated significant enrichment of genes related to cellular senescence (SAUL_SEN_MAYO) and SASP in 1.5 kPa (Figure 2E)^38,39^. Enzyme-linked immunosorbent assay (ELISA) confirmed increased secretion of IL8, a key SASP component, in cells cultured on 1.5 kPa substrates (Figure S2E).

Given the crucial role of p53 in regulating cellular senescence^40,41^, we examined whether senescence driven by soft substrates is mediated by p53. GSEA revealed enrichment of p53 pathway genes at 1.5 kPa (Figure S2F), indicating p53 activation. The p53 reporter assay validated these results (Figure 2F)^42^. Furthermore, p53 knockdown (KD) blocked cellular senescence induced by soft substrates (Figures 2G, S2G and S2H). Consistent with these findings, previous studies have demonstrated increased p53 expression in various senescent RPE models, including aged mice, with a corresponding reduction in p53 expression upon senescence alleviation^13,15^. These results suggest that diminished cell‒ECM interactions drive p53-dependent senescence in RPE cells.

### RPE senescence induced by declined integrin signaling

Integrins are crucial mediators in detecting and transmitting mechanical signals from the ECM to cells^30^. Given the significant role of ITGB1 in RPE cells, we employed shRNAs targeting ITGB1 (Figures S3A and S3B). ITGB1 depletion led to approximately 80-90% of ARPE19 cells being positive for SA-β-gal, along with an increased number of p21^+^ cells (Figures 3A and 3B). ITGB1 KD also increased the expression of SASP genes and reduced lamin B1 (*LMNB1*) expression, another hallmark of cellular senescence (Figure 3C)^43^. Additionally, blocking integrin from binding to the ECM using RGDS peptides increased the number of SA-β-gal^+^ senescent cells (Figures S3C and S3D), suggesting that perturbation of integrin-mediated cell‒ECM interaction induces RPE senescence.

**Fig. 3.**
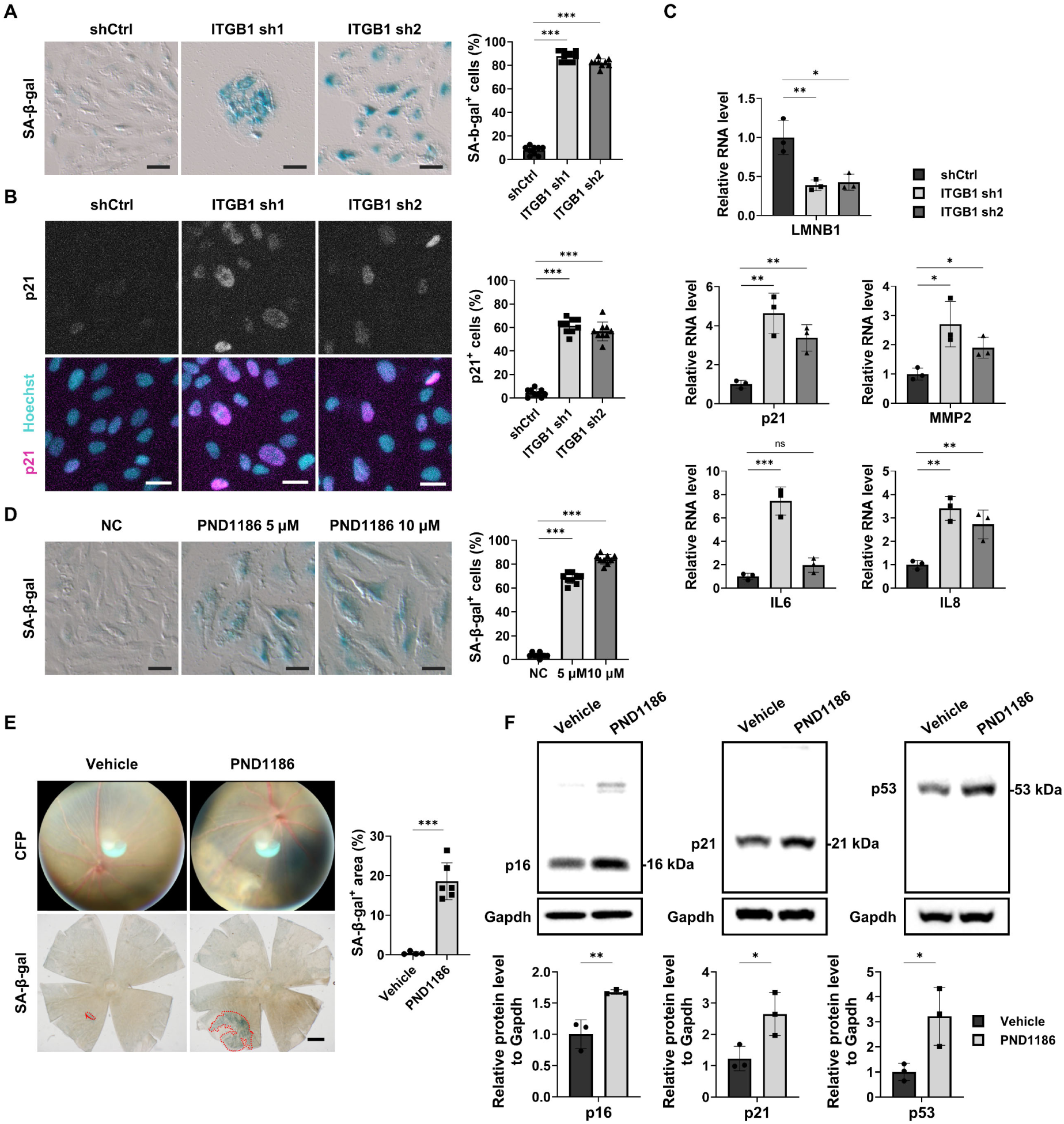
RPE senescence is induced by declined integrin signaling. (A and B) SA-β-gal staining (A), and immunofluorescence assay for p21 (B) in ARPE19 cells after ITGB1 KD. *n* = 9 regions from three independent experiments. (C) qPCR analysis of LMNB1, p21 and SASP genes in ARPE19 cells after ITGB1 KD. *n* = 3 samples from three independent experiments. (D) SA-β-gal staining in ARPE19 cells treated with a FAK inhibitor, PND1186 for 96 hours. *n* = 9 regions from three independent experiments. (E) Representative color fundus photographs (CFPs) (upper) and SA-β-gal staining of RPE/choroid flat mounts (lower) from C57BL/6J mice subretinally injected with PND1186 for 1 week. Red dashed lines in the RPE/choroid flat mounts highlight regions with elevated SA-β-gal activity. *n* = 4 mice for Vehicle and *n* = 6 mice for PND1186. (F) Western blot analysis of isolated RPE cells from mouse eyes for p16, p21, and p53 in PND1186-treated mice. *n* = 3 samples per group. Two-sided Student’s t-test, ***P < 0.001, **P < 0.01, *P < 0.05, ns; not significant. Exact *P* values are listed in Table S10. Scale bars: 25 μm (B), 50 μm (A, D), 500 μm (E)

FAK plays a pivotal role in integrin signaling, and its activity declined in the RPE cells of old mouse (Figures 1G and S1F). A reversible FAK inhibitor, PND1186, was used to investigate the contribution of FAK-mediated integrin signaling to cellular senescence (Figure S3E)^44^. FAK inhibition induced senescent phenotypes in ARPE19 cells with a high proportion of SA-β-gal^+^ and p21^+^ cells and transcriptional changes, including increased expression of SASP genes and decreased *LMNB1* expression (Figures 3D, S3F and S3G). To corroborate the role of FAK inhibition in RPE senescence *in vivo*, we administered PND1186 into the subretinal space of 8-week-old mice. Consistent with the *in vitro* findings, FAK inhibition significantly increased the SA-β-gal stained area, as well as the p16, p21, and p53 protein levels, in the young mouse RPE (Figures 3E and 3F). These results highlight the importance of dysregulated integrin signaling in driving RPE senescence.

### RPE senescence driven by YAP inactivation

YAP and TAZ (Transcriptional co-activator with PDZ-binding motif) are key transcription factors downstream of integrin signaling that facilitate the transmission of external mechanical cues to the nucleus^45–47^. Recent studies have highlighted the role of YAP in preventing cellular senescence in mesenchymal tissues and fibroblasts^48–50^. Conversely, another study posits the involvement of YAP in promoting the survival of senescent cells, indicating its context-dependent functions^51^. However, the regulation of YAP and its specific role in RPE aging has yet to be explored. To address this, we assessed YAP signature scores based on YAP target genes in the scRNA-seq data^52^. The young-dominant clusters exhibited significantly higher YAP scores than the old-dominant clusters (Figures 1C and 4A). Immunostaining revealed a marked reduction in nuclear Yap expression in aged RPE compared to that in young RPE (Figure 4B). In contrast, Yap expression in the photoreceptor layers was undetectable in both young and aged mice (Figure 4B). This finding suggests that age-related Yap regulation in the outer retina is predominantly localized in the RPE. Consistently, ARPE19 cells cultured on soft substrates displayed reduced YAP activity (Figures 4C, S4A and S4B). Inhibiting integrin signaling using PND1186 decreased the expression of well-established YAP target genes, such as CTGF and CYR61 (Figure S4C). These findings suggest that compromised integrin signaling during RPE aging contributes to YAP inactivation.

**Fig. 4.**
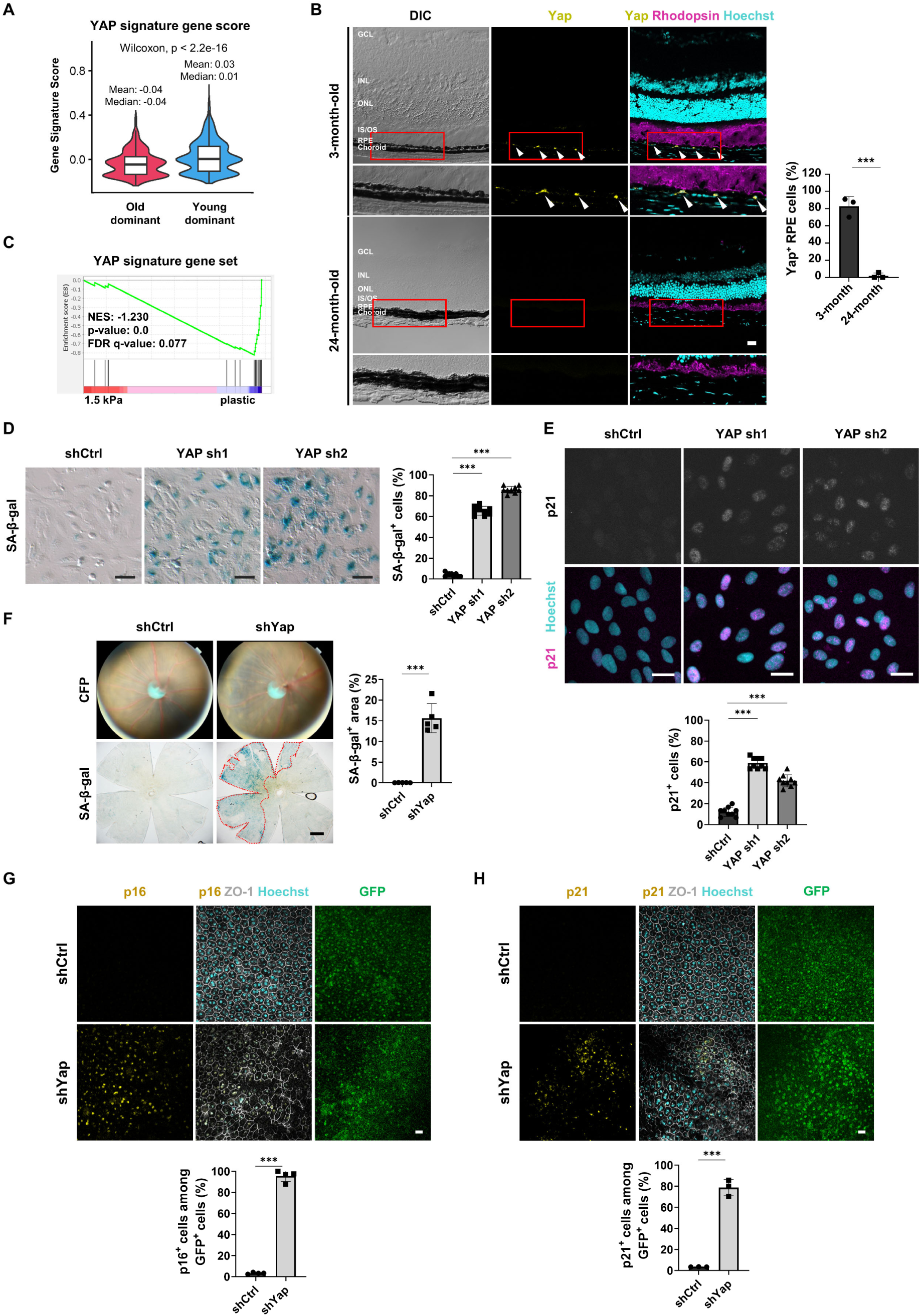
YAP inactivation drives RPE senescence. (A) Violin plot of the YAP signature gene score from the scRNA-seq data. (B) Immunostaining for YAP and rhodopsin (a marker for rod photoreceptors) in cryosectioned retina/RPE/choroid tissues from 3-month-old and 24-month-old mice. Arrowheads indicate YAP^+^ RPE cells in 3-month-old mice. YAP signal was minimal or undetectable in RPE cells from 24-month-old mice. Red boxes indicate the RPE region, which is shown at higher magnification in the lower panels. *n* = 3 samples per group. (C) GSEA analysis of the YAP signature gene set in the RNA-seq data from ARPE19 cultured either on plastic or 1.5 kPa substrate. (D and E) SA-β-gal staining (D), and immunofluorescence assay for p21 (E) in ARPE19 cells after YAP KD. *n* = 9 regions from three independent experiments. (F) Representative CFPs (upper) and SA-β-gal staining (lower) of RPE/choroid flat mounts from C57BL/6J mice subretinally injected with lentiviral vectors expressing Yap shRNA for 2 weeks. Red dashed lines in the RPE/choroid flat mounts highlight regions with elevated SA-β-gal activity. *n* = 5 mice per group. (G and H) Immunostaining for p16 (G), p21 (H), and ZO-1 in RPE cells infected with lentiviral vectors co-expressing Yap shRNA and GFP. *n* = 4 mice for p16 and *n* = 3 mice for p21. Abbreviations: GCL, ganglion cell layer; INL, inner nuclear layer; ONL, outer nuclear layer; IS, inner segment of photoreceptors; OS, outer segment of photoreceptors. Two-sided Student’s t-test, ***P < 0.001, **P < 0.01. Exact *P* values are listed in Table S10. Scale bars: 20 μm (B, G, H), 25 μm (E), 50 μm (D), 500 μm (F)

To explore whether YAP inactivation is sufficient to drive RPE senescence, we employed shRNAs targeting YAP (Figures S4D and S4E). YAP KD in ARPE19 cells markedly increased the number of SA-β-gal^+^ and p21^+^ cells, accompanied by elevated expression of SASP genes (Figures 4D, 4E and S4F). These results were corroborated using K-975^53^, a small molecule inhibitor that disrupts the protein‒protein interaction between YAP/TAZ and TEAD (Figures S4G and S4H). Furthermore, subretinal injection of shYap lentiviral vectors or K-975 in 8-week-old mice dramatically increased the number of senescent RPE cells marked by SA-β-gal, p16, and p21, highlighting the anti-senescent role of Yap *in vivo* (Figures 4F-H, S4I and S4J). These findings underscore that the age-related decline in YAP activity is instrumental in driving RPE senescence, suggesting that YAP activation could be a physiological approach to overcome RPE senescence driven by dysfunctional cell‒ECM interactions.

### Alleviating senescent RPE phenotypes through YAP activation

Given the pivotal role of YAP inactivation in RPE senescence, we investigated whether YAP activation could mitigate cellular senescence induced by impaired cell‒ECM adhesion. Ectopic YAP expression completely abrogated the emergence of SA-β-gal^+^ senescent cells on soft substrates (Figures 5A and S5A). Moreover, YAP overexpression rescued ITGB1 KD-induced senescence (Figure S5B). Bulk RNA-seq analysis was performed to comprehensively assess the transcriptomic effects of YAP overexpression in senescent ARPE19 cells on soft substrates (Figure 5B and Table S7). Upregulated DEGs in YAP-overexpressing cells (log_2_FC ≥ 1, adjP < 0.05) included cell cycle genes enriched in mitosis-related terms (Figure S5C). In contrast, downregulated DEGs (log_2_FC ≤ -1, adjP < 0.05) were enriched for terms related to innate immune responses (Figure S5D). Furthermore, genes associated with cellular senescence (SAUL_SEN_MAYO), p53 pathway, and SASP were significantly reduced in YAP-overexpressing cells compared to control senescent cells (Figure 5C). Given the role of YAP as a key stem cell factor, we examined the expression of a stemness-related gene set (RAMALHO_STEMNESS_UP) that is commonly expressed in embryonic, neural, and hematopoietic stem cells^54^. YAP overexpression activated this stem cell-related transcriptional program in senescent cells (Figure 5D). The overall transcriptomic profile of YAP-overexpressing cells on soft substrates clustered closely with that of cells cultured on plastic, highlighting the anti-senescence effect of YAP (Figure S5E).

**Fig. 5.**
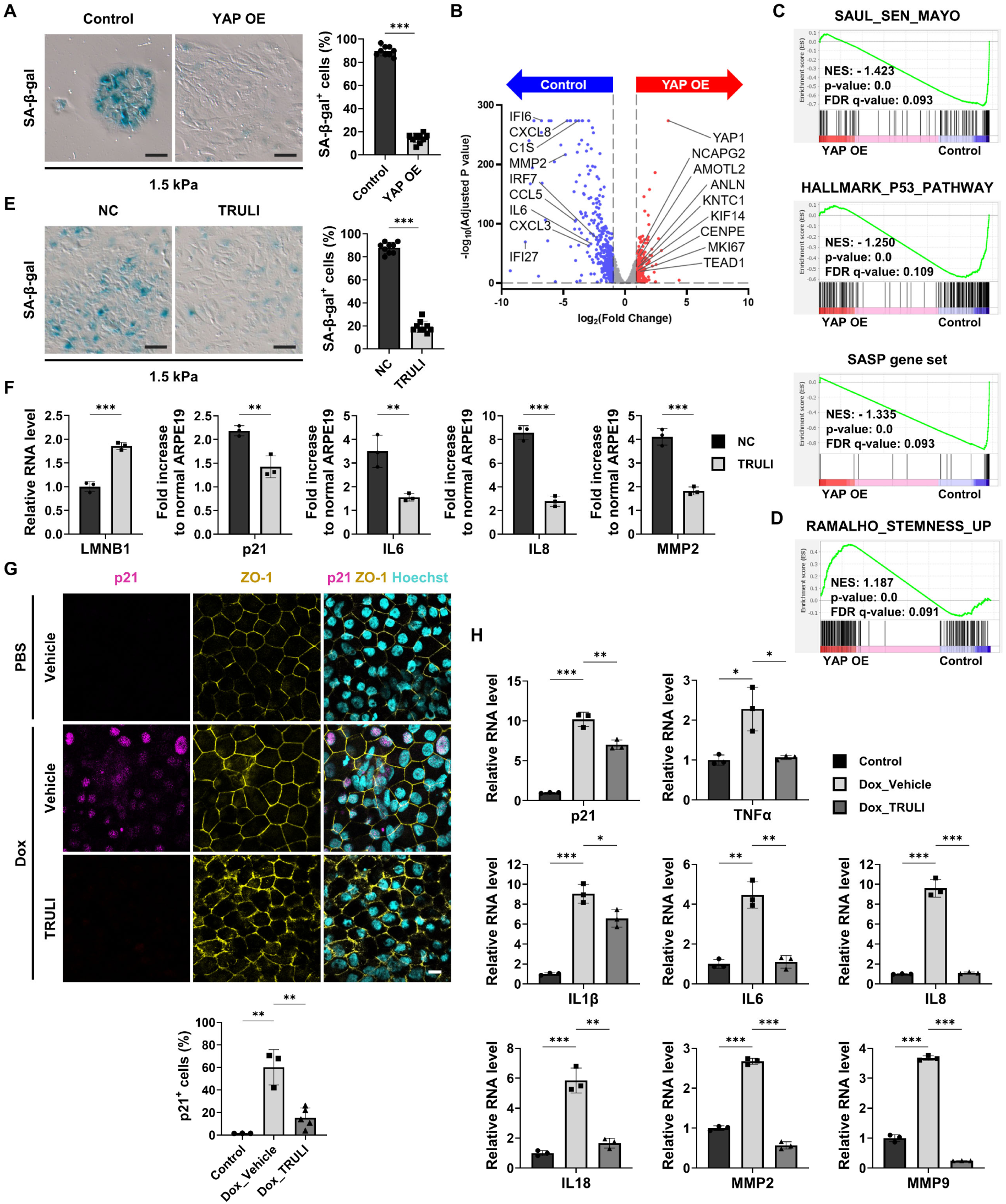
YAP activation alleviates senescent phenotypes of RPE. (A) SA-β-gal staining in ARPE19 cells cultured on 1.5 kPa after transduction with lentiviral vectors overexpressing YAP-HA. *n* = 9 regions from three independent experiments. (B) Volcano plot of DEGs from control and YAP-HA expressing ARPE19 cells cultured on 1.5 kPa substrate. The red and blue dots indicate either upregulated or downregulated genes after YAP overexpression, with cutoff values for DEGs: |log_2_FC| ≥ 1, adjP < 0.05, average TPM value ≥ 5. (C and D) GSEA with the RNA-seq data from (B). (E) SA-β-gal staining in ARPE19 cells cultured on 1.5 kPa and treated with TRULI (10 μM) for 24 hours. *n* = 9 regions from three independent experiments. (F) qPCR analysis of LMNB1, p21 and SASP genes in ARPE19 cells cultured on 1.5 kPa and treated with TRULI (10 μM) for 24 hours. *n* = 3 samples from three independent experiments. (G) Immunostaining for p21 and ZO-1 in hiPSC-derived RPE cells treated with Doxorubicin (250 nM) for 3 days, followed by a 24-hour treatment with TRULI (10 μM) on the 6th day. *N* = 3 for Control and Dox_Vehicle, and *n* = 5 for Dox_TRULI. (H) qPCR analysis of p21 and SASP gene mRNAs in Doxorubicin-exposed hiPSC-derived RPE cells treated with TRULI (10 μM). Two-sided Student’s t-test, ***P < 0.001, **P < 0.01, *P < 0.05. Exact *P* values are listed in Table S10. Scale bars: 10 μm (G), 50 μm (A, E)

We further investigated whether activation of endogenous YAP in already senescent cells could reverse the senescent phenotypes. TRULI, an ATP-competitive inhibitor of LATS1/2 kinases, was employed to activate YAP (Figures S5F and S5G). Treating ARPE19 cells cultured on soft substrates with TRULI reduced the number of SA-β-gal^+^ cells and decreased the expression of p21 and SASP genes, while restoring the expression of young cell marker *LMNB1* (Figures 5E and 5F). TRULI also exhibited anti-senescence activity in ARPE19 cells and human induced pluripotent stem cell (hiPSC)-derived RPE cells subjected to doxorubicin-induced senescence (Figures 5G, 5H and S5H-J). Propidium iodide (PI) staining confirmed that the anti-senescent effect of TRULI was not due to the selective death of senescent cells (Figure S5K). Previous studies have used complement-competent human serum (CC-HS) to recapitulate AMD cellular phenotypes in RPE cells^55^. CC-HS treatment significantly increased the expression of SASP genes in ARPE19 cells, which was mitigated by TRULI treatment (Figure S5L). These findings demonstrate that YAP activation, either through gene delivery or small molecules, alleviates senescent phenotypes in RPE cells.

### Restoration of visual function via Yap activation in an AMD-like mouse model

To assess the potential of TRULI in mitigating RPE senescence *in vivo* and its subsequent effect on retinal degeneration, we performed a subretinal injection of doxorubicin to induce RPE senescence. The presence of RPE cellular senescence in this model was previously validated using scRNA-seq analysis and key senescence markers^13–15,56^. TRULI was administered twice via intravitreal injection, and its efficacy in reducing RPE senescence was evaluated after 7 days (Figure S6A). Quantitative analysis of SA-β-gal staining in RPE/choroid flat mounts demonstrated a notable reduction in SA-β-gal^+^ area in the RPE layer of the TRULI-treated group compared to the vehicle control (Figure S6B). Additionally, immunofluorescence of RPE/choroid flat mounts revealed a decrease in the expression of the senescence markers p16 and p21, along with the restoration of the nuclear envelope protein Lmnb1 after TRULI treatment (Figures S6C-E). In doxorubicin-treated RPE cells, individual cell sizes increased and became more heterogeneous, with cells losing the uniform hexagonal shape characteristic of healthy RPE. TRULI treatment effectively restored cell morphology, leading to more uniform cell sizes and the re-establishment of the characteristic hexagonal structure (Figure S6E). Senescence reversal was further supported by structural improvements in both the RPE and retina, including reduced retinal degeneration in color fundus photography (CFP) and fundus autofluorescence (AF) imaging, as well as restoration of RPE layer integrity and outer nuclear layer (ONL) thickness (Figure S6F). Electroretinogram (ERG) recordings demonstrated a significant recovery in both c- and a/b-wave amplitudes, indicating enhanced RPE and retinal function, respectively, in TRULI-treated mice (Figure S6G and S6H).

Next, the effect of TRULI on RPE senescence was investigated in an AMD-like mouse model induced by sodium iodate (NaIO_3_), a well-characterized late-stage dry AMD or GA model. Retinal degeneration is triggered by oxidative damage to the RPE, leading to secondary photoreceptor degeneration^57–59^. Because RPE senescence has not been previously reported in this model, we investigated the onset of senescence prior to severe RPE damage. Mice received a 25 mg/kg NaIO_3_ intraperitoneal injection, and RPE senescence was analyzed through SA-β-gal and p16/p21 staining at days 0, 1, 3, and 7, alongside cleaved caspase-3 staining (Figure S7A). RPE senescence was detectable by day 3 and pronounced by day 7 (Figures S7B-E), whereas no evidence of retinal senescence was observed during the 7-day period (Figure S7C). Importantly, RPE senescence preceded apoptosis and retinal degeneration, which became evident only on day 7 (Figures S7B and S7F). This temporal pattern demonstrates that the model replicates the progression of RPE senescence and its role in initiating retinal degeneration.

TRULI treatments, administered intravitreally over four intervals starting 3 days post-NaIO_3_ injection to male mice (Figure 6A), significantly reduced senescent RPE areas (Figure 6B). Immunofluorescence and mRNA analyses confirmed reduced p16 and p21 expression in the TRULI-treated group (Figures 6C-E). TRULI treatment restored cell morphology and nuclear envelope integrity, as evidenced by the recovery of Lmnb1 expression around RPE nuclei (Figure 6F). Anatomical improvements in the retinas of TRULI-treated mice were evident with CFP imaging and hematoxylin and eosin (H&E) staining, demonstrating recovery of the RPE layer and ONL thickness (Figure 6G). AF imaging revealed a reduction in hyperautofluorescent spots, suggesting the role of TRULI in reducing the number of senescent metabolically active RPE cells (Figure 6G). Functional improvements were confirmed by ERG, in which the treated mice showed significant enhancements in both c- and a/b-wave responses, reflecting restored visual function (Figure 6H and 6I). Similar anti-senescent effects and morphological improvements were also observed in female mice treated with TRULI in the NaIO₃ model (Figures S8A-C), suggesting that TRULI’s therapeutic effects are not sex-dependent. Furthermore, these beneficial effects persisted for up to two months post-treatment, as evidenced by sustained reduction in p16 and p21 expression and preservation of RPE structure (Figures S8A-C).

**Fig. 6.**
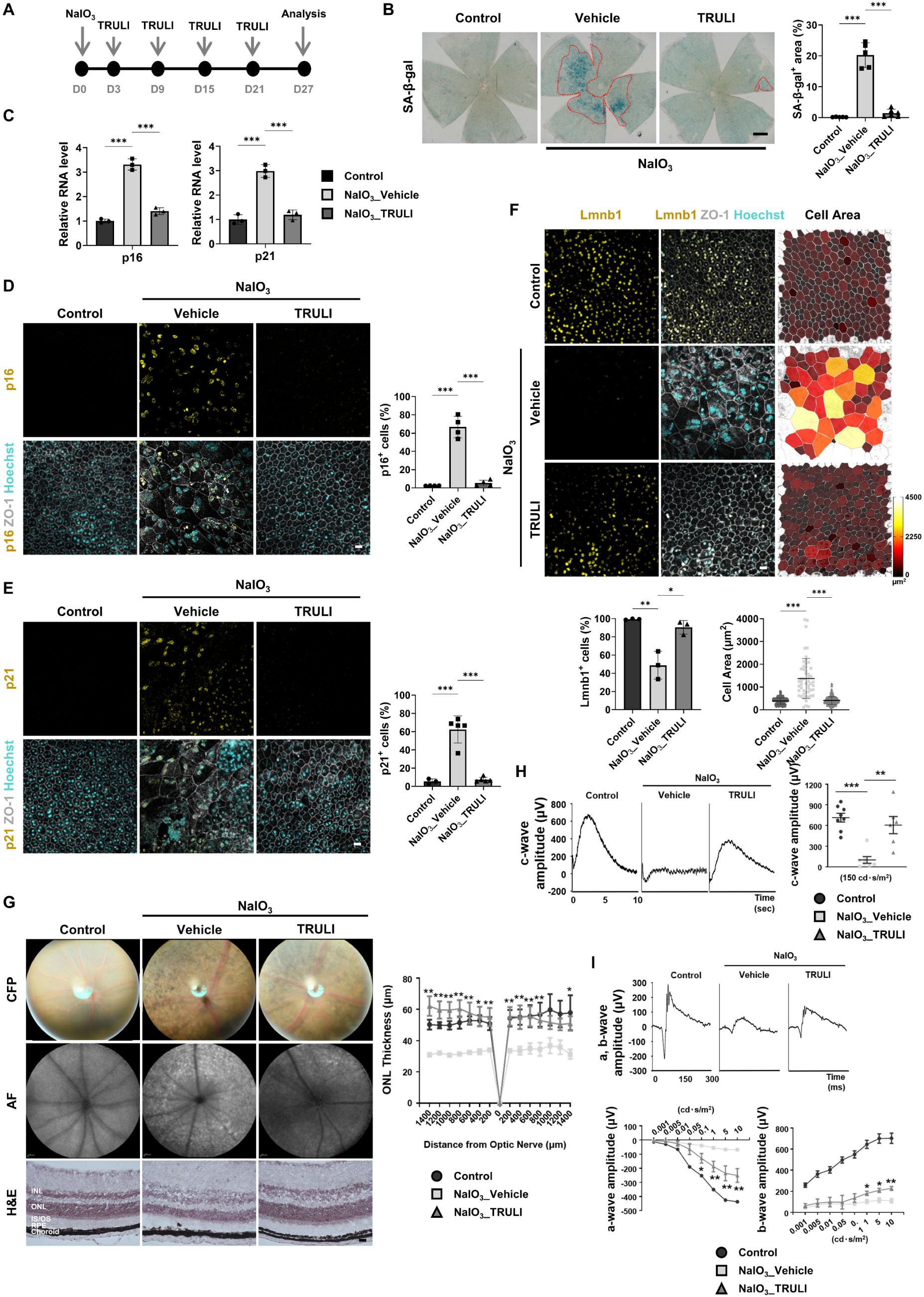
Yap activation in an AMD-like mouse model restores visual function. (A) Experimental scheme for evaluating TRULI effects on visual function in the AMD-like pathology mouse model. (B) SA-β-gal staining in RPE/choroid flat mounts. Red dashed lines in the RPE/choroid flat mounts highlight regions with elevated SA-β-gal activity. *n* = 5 mice for control and NaIO_3__Vehicle, and n = 6 mice for NaIO_3__TRULI. (C) qPCR analysis of p16 and p21 in RPE from the AMD-like pathology mouse model injected with TRULI. *n* = 3 samples from three independent experiments. (D, E, and F) Immunostaining for p16 (D), p21 (E), Lmnb1 (F), and ZO-1 in RPE/choroid flat mounts. n = 4 mice per group in (D); n = 3 mice for control, n = 5 mice for NaIO_3__Vehicle and NaIO_3__TRULI in (E); *n* = 3 mice per group in (F). The measured cell areas (F) were visualized using color mapping with the LUT Red Hot scale. *n* = 229 cells for control, *n* = 52 for NaIO_3__Vehicle, and n = 217 for NaIO_3__TRULI. (G) Representative CFPs (upper), autofluorescence (AF) images (middle), and H&E-stained retinas (lower) are shown. The thickness of the outer nuclear layer (ONL) was measured at 200 μm intervals from the center of the optic nerve. *n* = 5 mice per group. (H and I) ERG analysis in NaIO_3_-induced mice treated with TRULI. The amplitude of the c-wave (μV) was measured at 150 cd·s/m² (H). *n* = 8 mice for control, *n* = 7 mice for NaIO_3__Vehicle, and *n* = 6 mice for NaIO_3__TRULI. The amplitudes of the scotopic a-wave and b-wave were measured across a range of 0.001-10 cd·s/m² (I). *n* = 6 mice for control and NaIO_3__Vehicle, *n* = 4 mice for NaIO_3__TRULI (a-wave). n = 6 mice for control, n = 4 mice for NaIO_3__Vehicle and NaIO_3__TRULI (b-wave). Two-sided Student’s t-test, ***P < 0.001, **P < 0.01, *P < 0.05. Exact *P* values are listed in Table S10. Scale bars: 20 μm (D, E, F, G), 500 μm (B)

The intravitreal injection of TRULI can affect all retinal cell types. An additional experiment using subretinal injection of AAV-Yap was performed to determine whether YAP activation in the RPE plays a role in restoring visual function in NaIO_3_-treated mice. Low concentrations of AAV2/8 are known to preferentially target the RPE following subretinal delivery^60,61^. RPE-specific delivery of AAV was confirmed in mice and predominant mCherry signals were observed in the RPE with minimal expression in the retina (Figures S9A and S9B). RPE-targeted Yap overexpression markedly reduced senescent RPE areas, decreased p16 and p21 expression, restored Lmnb1 levels, and recovered uniform cell morphology in male mice two weeks after AAV-Yap injection (Figures S9C-F). These cellular improvements were accompanied by anatomical and functional restoration of the RPE and retina, as demonstrated by CFP, H&E staining, and enhanced ERG responses in NaIO_3_-treated mice (Figures S9G-I). Similarly, AAV-Yap administration in female mice also led to significant reductions in RPE senescence markers and structural restoration at both two weeks and two months post-injection (Figure S10A-C), indicating durable therapeutic effects of YAP activation. Collectively, these findings indicate that Yap activation in this AMD-like mouse model reverses RPE senescence, thereby restoring visual function.

### RPE rejuvenation in naturally aged mice through Yap activation

To investigate RPE rejuvenation in naturally aged mice, mice aged 23 months were treated with TRULI for 4 weeks (Figure 7A). TRULI treatment significantly reduced the number of SA-β-gal^+^ RPE cells, decreased p16 and p21 expression, and restored RPE morphology with uniform monolayer architecture, leading to more consistent cell size and structure in RPE/choroid flat mounts (Figures 7B-D). Furthermore, diminished Lmnb1 immunofluorescence in the RPE nuclei of aged mice was restored to levels comparable to those in younger controls (Figure 7E). Corresponding CFP images revealed fewer atrophic patches, and H&E staining demonstrated the recovery of ONL thickness in TRULI-treated mice (Figure 7F). Whole transcriptome analysis of RPE cells revealed that downregulated DEGs (log_2_FC ≤ -1, adjP < 0.05) in TRULI-treated mice were enriched in terms related to immune and defense response (Figures 7G, 7H and Table S8). In contrast, upregulated DEGs (log_2_FC ≥ 1, adjP < 0.05) were highly enriched in embryonic developmental terms, demonstrating the role of Yap in activating stem cell transcriptional programs (Figure 7I). ERG recordings showed substantial improvements in both the c- and a/b-wave responses in TRULI-treated mice (Figures 7J and 7K). These findings suggest that TRULI-induced Yap activation increases the expression of developmental genes and alleviates RPE senescence, leading to substantial improvements in visual function in naturally aged mice.

**Fig. 7.**
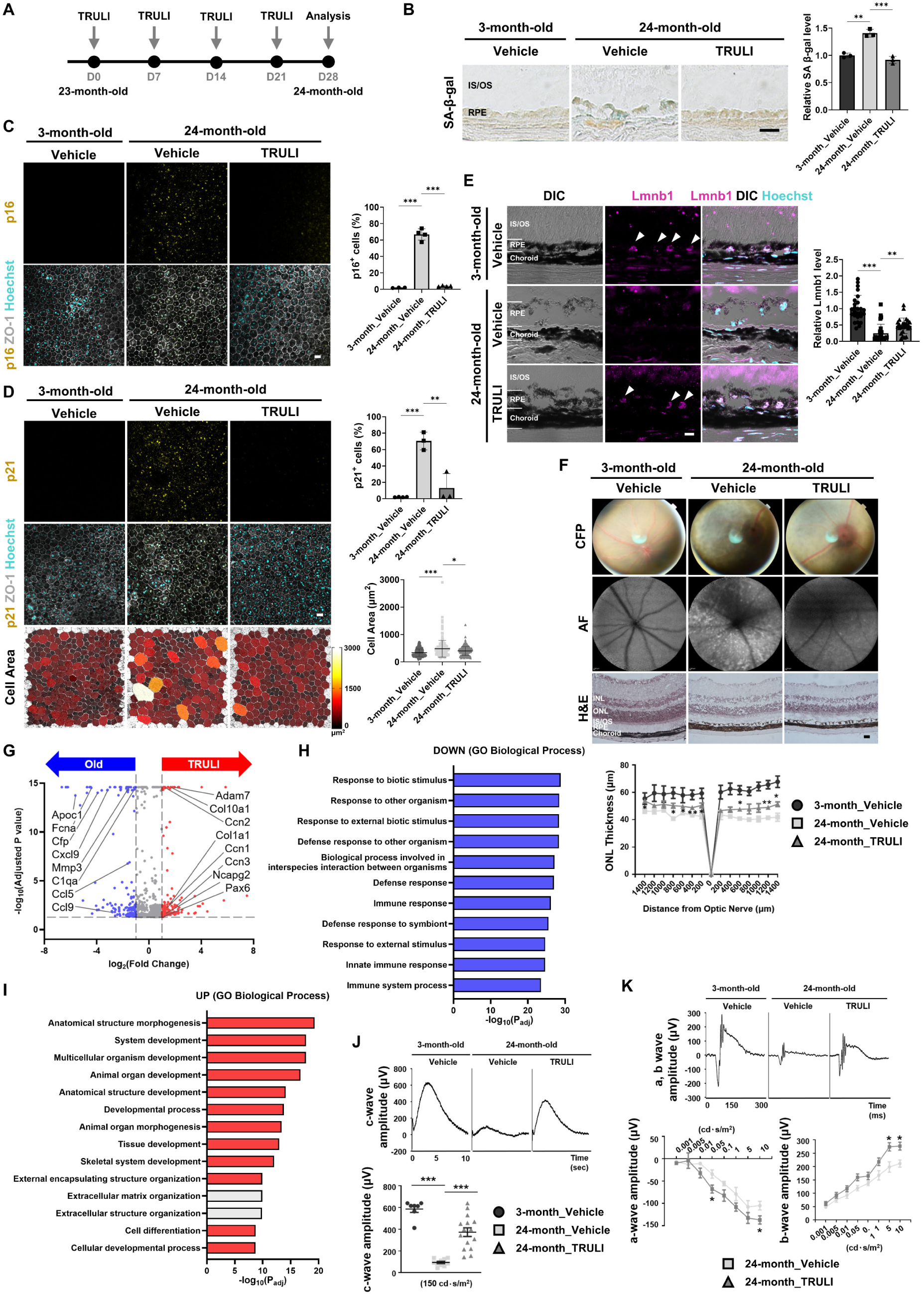
Yap activation by TRULI rejuvenates visual function in aged mice. (A) Experimental scheme for evaluating TRULI effects on visual function in aged mice. (B) SA-β-gal staining in cryosectioned retina/RPE/choroid. *n* = 3 samples per group. (C and D) Immunostaining for p16 (C), p21 (D), and ZO-1 in RPE/choroid flat mounts. *n* = 3 mice for 3-month_Vehicle, *n* = 4 mice for 24-month_Vehicle and 24-month_TRULI in (C); *n* = 4 mice for 3-month_Vehicle, n = 3 mice for 24-month_Vehicle and 24-month_TRULI in (D). The measured cell areas were visualized using color mapping with the LUT Red Hot scale. n = 259 cells for 3-month_Vehicle, n = 176 cells for 24-month_Vehicle, n = 215 cells for 24-month _TRULI in (D). (E) Immunostaining for Lmnb1 in cryosectioned retina/RPE/choroid. Arrowheads indicate Lmnb1^+^ RPE cells. Quantification of relative Lmnb1levels was performed in 26 RPE cells from 3 mice per group. (F) Representative CFPs (upper), AF images (middle), and H&E-stained retinas (lower). ONL thickness was measured at 200 μm intervals from the optic nerve center. *n* = 3 mice per group. (G) Volcano plot of DEGs from RPE of control or TRULI-treated aged mice. The red and blue dots indicate either upregulated or downregulated genes in TRULI-treated RPE, with cutoff values for DEGs: |log_2_FC| ≥ 1, adjP < 0.05, average TPM value ≥ 5. (H and I) The top pathways ranked by adjusted p-values from GO analysis with downregulated (H) and upregulated (I) genes from (G). (J and K) ERG analysis in 24-month-old mice treated with TRULI. The amplitude of the c-wave (μV) was measured at 150 cd·s/m² (J). *n* = 7 mice for 3-month_Vehicle, n = 10 mice for 24-month_Vehicle, *n* = 15 mice for 24-month_TRULI. The amplitude of the scotopic a-wave and b-wave was measured across the range of 0.001-10 cd·s/m² (K). *n* = 3 mice for 24-month_Vehicle, *n* = 5 mice for 24-month_TRULI. Two-sided Student’s t-test, ***P < 0.001, **P < 0.01, *P < 0.05. Exact *P* values are listed in Table S10. Scale bars: 20 μm (B, C, D, E, F)

## DISSCUSION

Current treatments for AMD primarily address specific pathological symptoms, with limited potential to halt or reverse disease progression^62,63^, highlighting the urgent need for safe and effective therapies targeting the root mechanisms of AMD. In this study, scRNA-seq analysis of the RPE from young and old mice identified disrupted cell‒ECM interactions as a hallmark of RPE senescence. Furthermore, the integrin-YAP mechanotransduction pathway was crucial for regulating RPE senescence *in vitro* and *in vivo*. YAP activation reversed RPE senescence and restored visual function in both AMD and naturally aged mouse models. These findings suggest that senescent RPE phenotypes are reversible and offer a promising therapeutic approach for AMD.

GWAS have identified SNPs linked to AMD^64^, many of which are associated with genes involved in ECM turnover and remodeling, including *HTRA1, TIMP3, ADAMTS9,* and *COL8A1*. *HTRA1* encodes a serine protease that cleaves ECM proteins such as fibronectin, whereas *ADAMTS9* encodes a metallopeptidase that degrades proteoglycans such as aggrecan and versican. TIMP3 inhibits metallopeptidase activity, and COL8A1, localizes to BrM, plays a key role in maintaining structural integrity. These AMD-associated SNPs suggest a strong link between AMD and cell‒ECM interactions, supporting the finding that impaired cell‒ECM adhesion is a crucial driver of RPE senescence and age-related visual dysfunction.

Integrins, a family of transmembrane receptors, mediate cell‒ECM interactions, including RPE cell adhesion to BrM. Integrins transduce mechanical cues from the ECM into intracellular signaling pathways and are central to mechanotransduction, thereby regulating cellular processes such as cell migration, proliferation, and survival. Integrins have been implicated in both promoting and protecting against cellular senescence; however, their precise role in senescence remains unclear. Integrin signaling contributes to cellular senescence by activating pathways like reactive oxygen species (ROS), PI3K/Akt, NF-κB, and TGF-β^65–69^. On the other hand, recent studies suggest that integrins also play a protective role against cellular senescence^70^. Variability in their effects may stem from differences in the mechanical properties of the microenvironments across tissues. In the case of RPE cells, they adhere to BrM, which has a very stiff mechanical property (7–19 MPa)^34^. Consistently, we observed strong integrin activity in the young RPE cells (Figure 1G). However, age-related drusen accumulation disrupts tight integrin‒BrM interactions and promotes RPE senescence. These findings emphasize the tissue-specific role of integrins in regulating cellular senescence.

The ectopic expression of pluripotency factors (OSKM) reprograms somatic cells into pluripotent stem cells through a process known as cellular reprogramming^71,72^. Partial reprogramming does not alter cell identity but reverses aging markers, enhances tissue repair in aged mice, and extends the lifespan of progeroid mice^21–23^. Recently, a cocktail of six chemicals used in chemically induced reprogramming reversed transcriptomic age^73^. However, the clinical application of multiple reprogramming factors remains challenging. In contrast, YAP activation alone can reprogram differentiated cells into tissue-specific stem cells^20^. Similar to OSKM, YAP overexpression reactivates the stem cell transcriptional program, reversing the senescent state of RPE cells. Furthermore, TRULI, a small-molecule YAP activator, significantly restored visual function in an AMD-like mouse model and naturally aged mice. Whole-transcriptome analysis confirmed the efficacy of TRULI in reversing age-related gene expression profiles. Unlike OSKM, which is not physiologically expressed in RPE, YAP is endogenously expressed in normal, young RPE cells and plays a crucial role in preventing cellular senescence. While the primary experiments in this study were conducted in male mice, subsequent validation in age-matched female mice demonstrated comparable anti-senescent effects and structural improvements following YAP activation. Given that AMD affects both sexes, these results further underscore the broad translational relevance of this approach. Thus, YAP activation represents a simpler, more physiologically relevant, and clinically feasible approach for reversing RPE aging and improving visual function.

Similar to other reprogramming factors, YAP is involved in oncogenic processes^74,75^. Therefore, mitigating its oncogenic potential is essential for successful rejuvenation treatments. The retina is an ideal target for testing the rejuvenating role of YAP owing to its relative isolation and the extremely low incidence of RPE or retinal tumors^76–78^. Thus, our findings provide a compelling example of YAP activation as a potential approach to address age-related diseases.

## RESOURCE AVAILABILITY

### Lead contact

Further information and requests for resources and reagents should be directed to and will be fulfilled by the lead contact, Jiwon Jang.

### Materials availability

Plasmids and other materials generated in this study are available from the lead contact upon request.

### Data and code availability

scRNA-seq and bulk RNA-seq data from this study are publicly accessible in the Gene Expression Omnibus (GEO) data repository at the National Center for Biotechnology Information (NCBI) under accession number GSE282283. The published single-cell RNA-seq data that were re-analyzed here are available from the Gene Expression Omnibus (GSE210543 and GSE203499).

## ACKNOWLEDGEMENTS

We thank Dr. Min Jae Song at NCATS/NIH and Dr. Kapil Bharti at NEI/NIH for insightful comments. This work was supported by the National Research Foundation of Korea (NRF) grants funded by the Government of Korea (Ministry of Science and ICT) (RS-2020-NR046273 to JJ; RS-2024-00334730 to JJ; RS-2025-00553501 to HC; RS-2021-NR058886 to SL; RS-2018-NR031072 to SL), and by a grant of the Korean ARPA-H Project through the Korea Health Industry Development Institute (KHIDI), funded by the Ministry of Health & Welfare, Republic of Korea (RS-2025-25454860 to HC).

## AUTHOR CONTRIBUTIONS

Conceptualization, J.J. and H.C.; methodology, G.K. and C.S.; validation, S.O., J-B.C., C-W.P.; formal analysis, H.K.L., J.C., S.L., and S.L.; investigation, G.K., C.S., H.K.L., S.O., J-B.C., and C-W.P.; writing – original draft, G.K., J.J., and H.C.; writing – review and editing, G.K., C.S., H.K.L., S.O., J-B.C., C-W.P., C.K., J-H.R., S.L., J.J., and H.C.; supervision and funding acquisition, S.L., J.J., and H.C.

## STAR★METHODS

### KEY RESOURCES TABLE

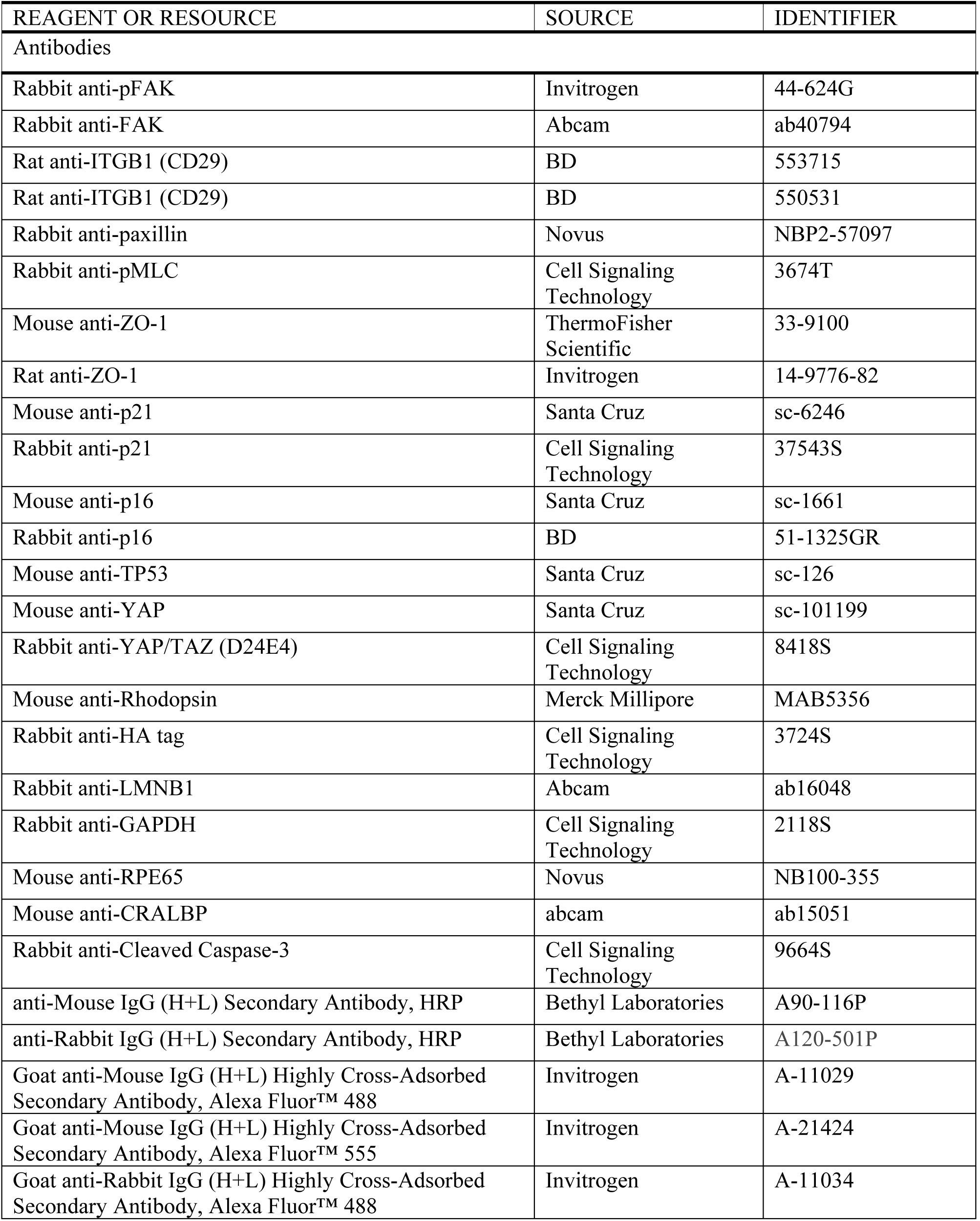
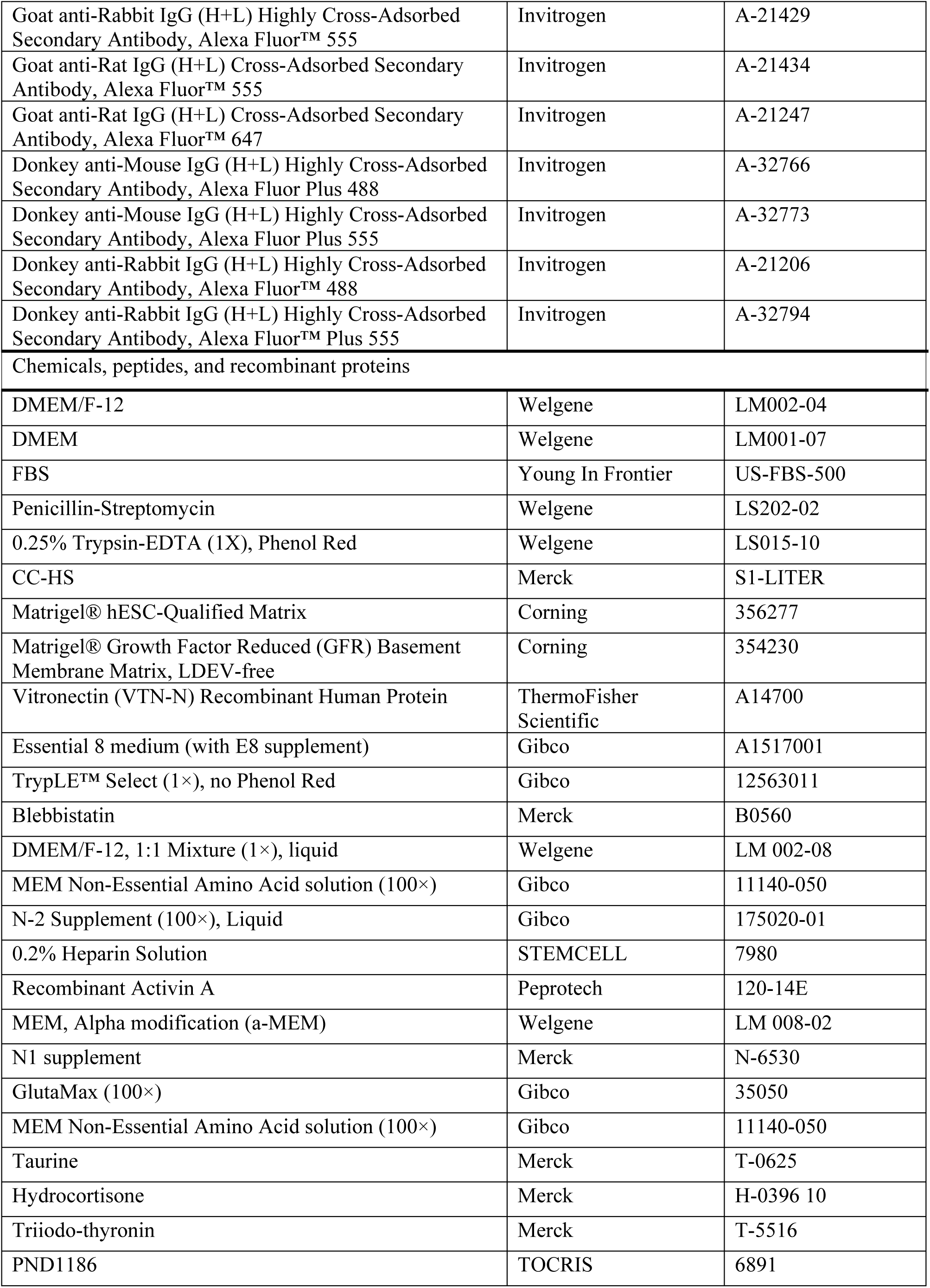
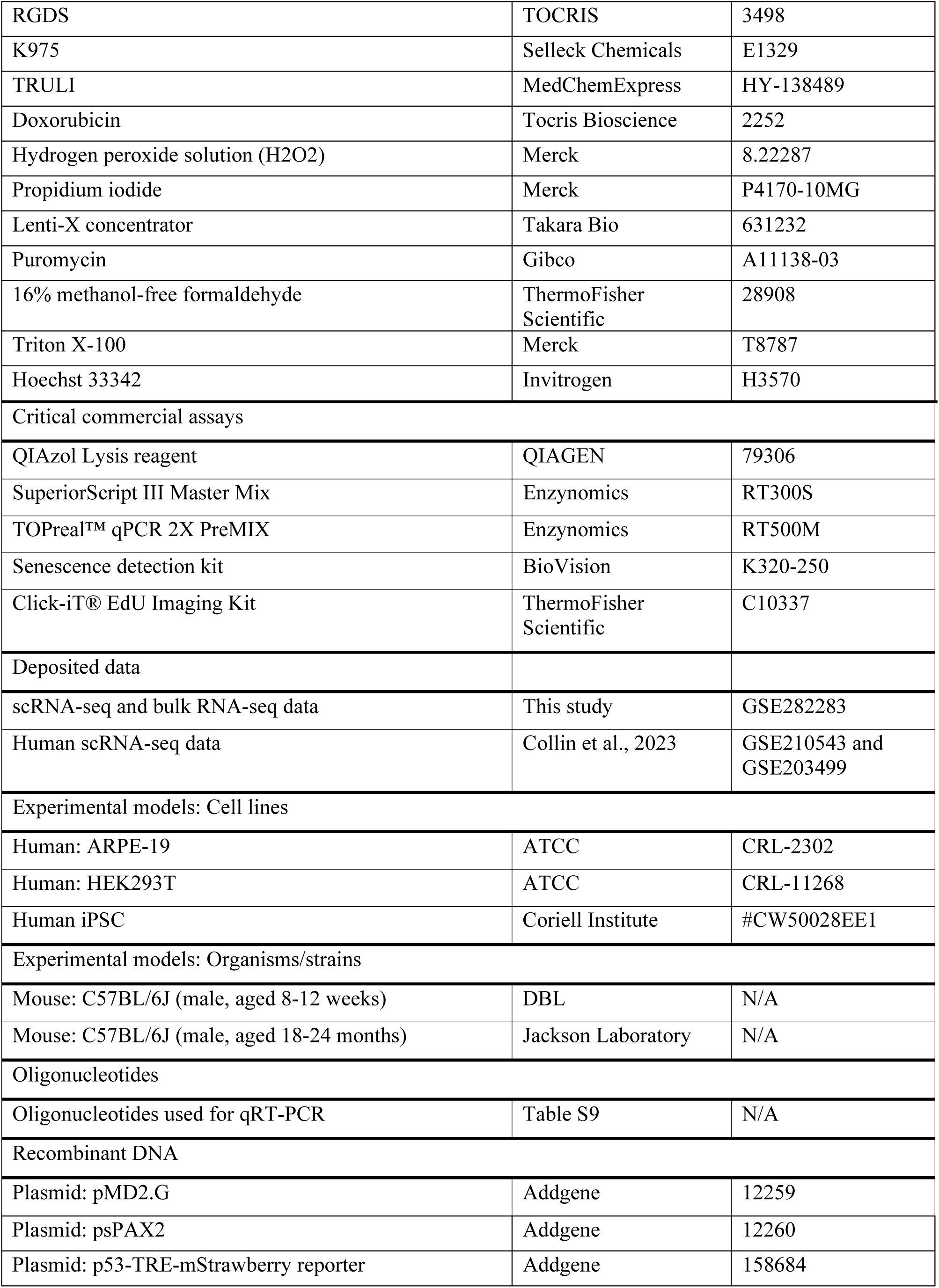
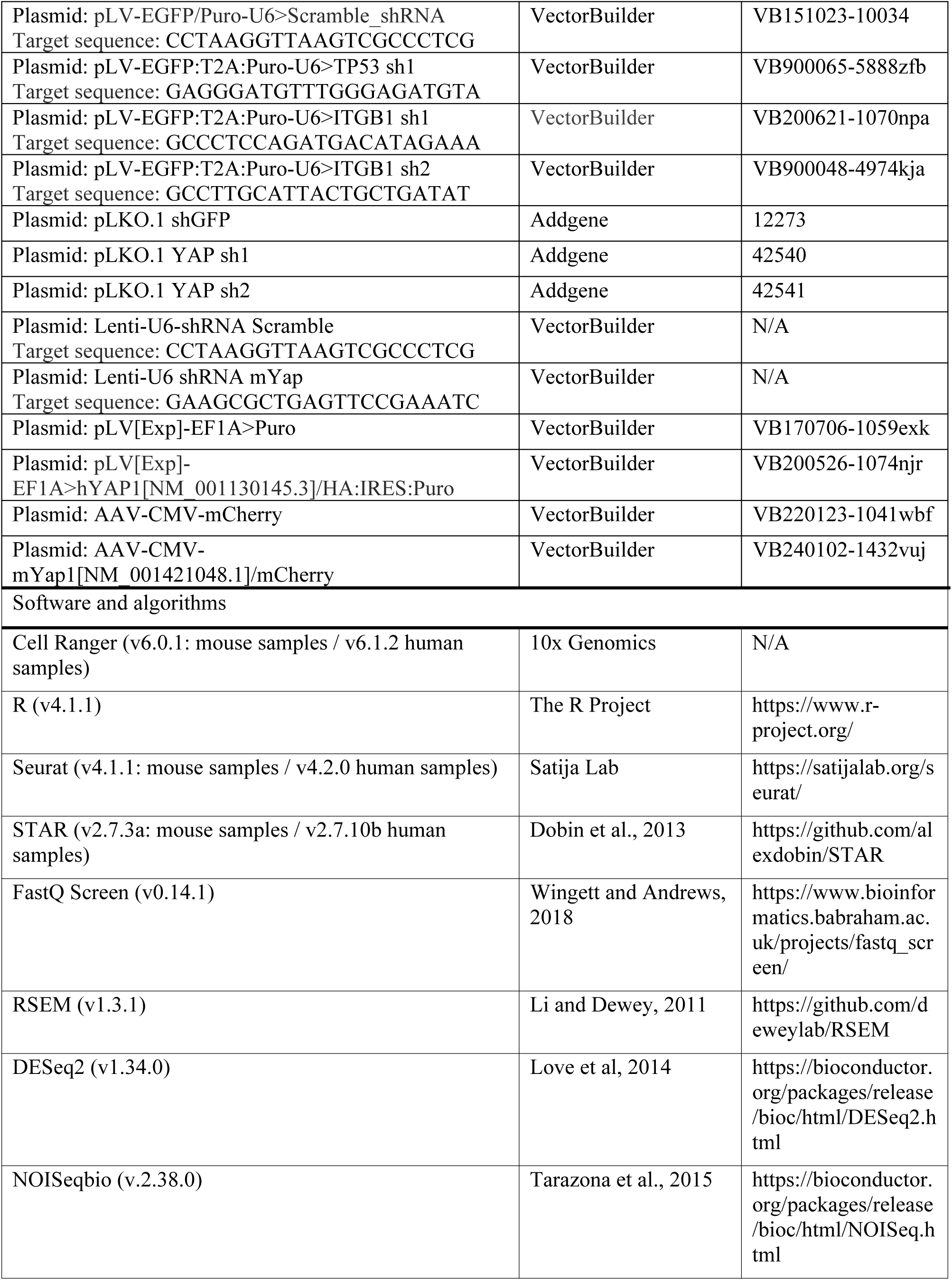
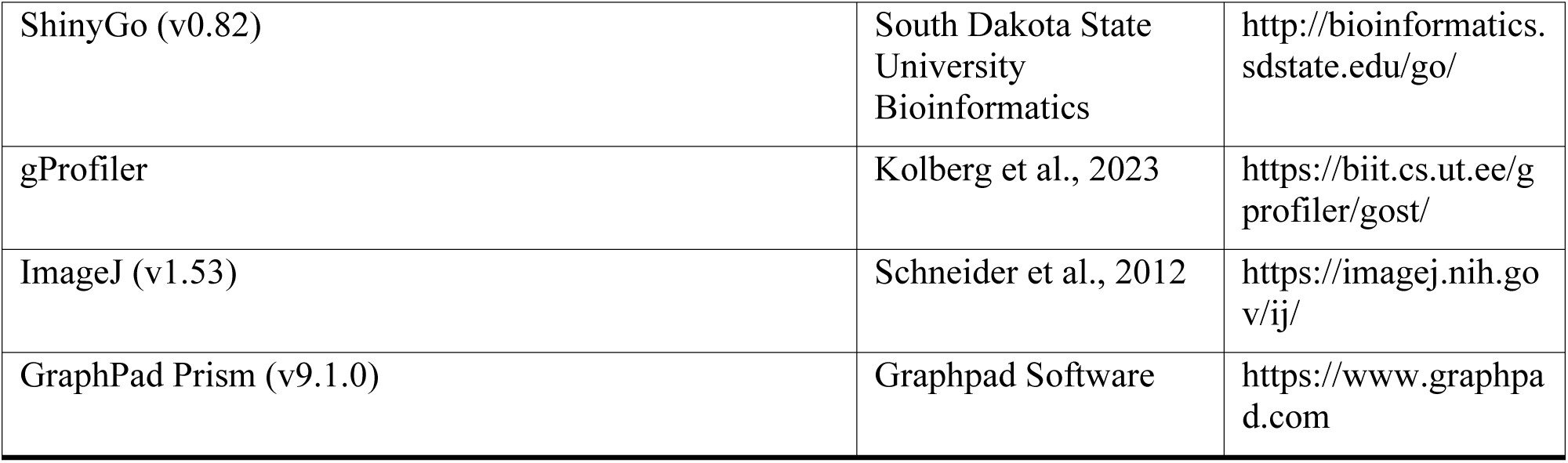

## EXPERIMENTAL MODEL AND SUBJECT DETAILS

ARPE19 cells (ATCC, CRL-2302) were maintained in DMEM/F-12, and HEK293T cells (ATCC, CRL-11268) were maintained in DMEM, all of which were supplemented with 10% fetal bovine serum (FBS) and 1% penicillin-streptomycin (Welgene). Cells were cultured at 37°C with 5% CO_2_ and passaged using trypsin (Welgene) every 2-3 days. All cell lines were routinely tested for mycoplasma contamination and confirmed to be negative.

All animal experiments were conducted in accordance with the guidelines approved by the Institutional Animal Care and Use Committee (IACUC) at Konkuk University (approval no. KU23146). The mice were housed under a 12-hour light/dark cycle with ad libitum access to food and water. 8-week-old C57BL/6J mice were obtained from DBL (Incheon, Korea), and 18-month-old mice were purchased from Jackson Laboratory (Bar Harbor, Maine, USA). The 18-month-old mice were maintained until 24 months of age at the Konkuk University Laboratory Animal Research Center under specific pathogen-free conditions. Unless otherwise specified, all mice were male C57BL/6J mice and were randomly allocated to experimental groups. For selected validation studies involving NaIO₃-induced degeneration and treatment of TRULI or AAV-Yap, age-matched female C57BL/6J mice were used following the same experimental protocols.

## METHOD DETAILS

### Generation of senescence models

To manipulate stiffness, ARPE19 cells were plated and cultured on polydimethylsiloxane gels with stiffnesses of 28 and 1.5 kPa (ibidi). Before plating, a thin Matrigel coating was applied to enable cell attachment. To establish a DNA damage-induced senescence model, ARPE19 cells were treated daily with 250 nM doxorubicin for 72 h. Doxorubicin was withdrawn and replaced with fresh medium every other day for 96 h, followed by TRULI treatment. Cell death was detected using PI staining (10 μg/mL). ARPE19 cells treated with 2 mM H_2_O_2_ (Sigma) for 12 h were used as positive controls. To induce AMD cellular phenotypes, ARPE19 cells were treated with 5% CC-HS (Millipore) in a complete medium for 5 days, followed by TRULI treatment in a medium containing 5% CC-HS.

### Differentiation of RPE from hiPSCs

hiPSCs were obtained from the Coriell Institute Biobank (Coriell, #CW50028EE1) and cultured in a feeder-free system on vitronectin-coated plates (Thermo). The protocol for inducing early retinal and subsequent RPE differentiation was adapted from previously published methods^79–82^.

To initiate neural progenitor differentiation, hiPSCs were enzymatically detached using TrypLE Select Enzyme (Thermo) and cultured as aggregates in suspension within a neural induction medium (NIM) for 7 days. The NIM formulation included DMEM/F-12, 1% N-2 supplement, MEM non-essential amino acids, and 2 µg/mL heparin. On day 7, the resulting embryoid bodies were transferred to Matrigel-coated plates (Corning) and cultured in RPE differentiation medium (RPE-DM), which comprised alpha-MEM (Welgene), 1× N1 supplement (Sigma), taurine (Sigma), triiodothyronine (Sigma), hydrocortisone (Sigma), non-essential amino acids (Thermo), penicillin-streptomycin (Welgene), and 5%–15% FBS (Thermo). To promote RPE differentiation, 10 ng/mL of activin A (Peprotech) was added to the culture medium for 14 days, with medium changes every 2 days. By day 50, RPE cells had fully matured and formed a confluent monolayer of pigmented cells. Doxorubicin-induced senescence and TRULI administration were performed in the same manner as described for ARPE19 cells.

### Lentivirus packaging and application

Lentivirus production was performed by co-transfecting HEK293T cells with 2^nd^ generation packaging vectors, psPAX2 (Addgene), pMD2.G (Addgene), and lentiviral vector plasmids using Lipofectamine 3000 (Invitrogen). The lentivirus-containing supernatant was collected and filtered through 0.45 μm cellulose acetate membrane filters (Corning) 48 h after transfection. The viral supernatant was then concentrated using a Lenti-X concentrator (Takara Bio) at a 3:1 ratio and incubated at 4°C for 30 min to overnight. The mixture was then centrifuged at 1,500 × *g* for 45 min at 4°C to pellet the virus. The virus pellet was resuspended in phosphate-buffered saline (PBS) and stored at -80°C. Cells were then transduced with pellet aliquots and selected with puromycin (1 μg/mL) for at least 48 h post-transduction. Lentiviruses, each at a concentration of 1 × 10⁷ TU/μL, were subretinally injected into C57BL/6J mice for transduction. The mice were euthanized two weeks later for analysis.

### SA-β-gal and EdU incorporation assays

The samples were fixed with a fixative solution (BioVision) at room temperature (RT) for 10 min. After fixation, the samples were washed with PBS and incubated overnight at 37°C in the SA-β-gal staining solution, prepared following the manufacturer’s instructions. After staining, the samples were washed three times.

Click-iT EdU imaging (Thermo Fisher Scientific) and senescence detection kits were used on the same samples to label the proliferating and senescent cells simultaneously. Before fixation, ARPE19 cells were treated with EdU at a final concentration of 10 μM at 37°C for 24 h. Cells were fixed using a fixative solution from the senescence detection kit, and SA-β-gal staining was performed as previously described. After SA-β-gal staining, samples were permeabilized, and EdU was detected following the Click-iT reaction protocol. Images were captured with a fluorescence microscope (Leica MICA), prepared, and quantified using the ImageJ 1.53 software (Fiji). Statistical data were analyzed and plotted using the GraphPad Prism 10.4.1 software (GraphPad Software).

### ELISA

The cell culture supernatant was obtained from the harvested culture medium and centrifuged at 1,000 × *g* for 10 min. Total IL-8 levels were detected using a Human IL-8 ELISA kit (Boster Bio) and normalized to the cell number. Statistical data were analyzed and plotted using the GraphPad Prism 10.4.1 software (GraphPad Software).

### Quantitative real-time PCR

Total RNA was isolated from the samples using QIAzol Lysis Reagent (QIAGEN) and reverse transcribed into complementary DNA using SuperiorScript III Master Mix (Enzynomics). Quantitative RT-PCR analysis was performed on the CFX Connect Real-Time PCR Detection System (Bio-Rad) with TOPreal™ qPCR 2X PreMIX (Enzynomics). For normalization, GAPDH was used as a control for all samples. The primers used are listed in Table S9. Statistical data were analyzed and plotted using the GraphPad Prism 10.4.1 software (GraphPad Software).

### Intravitreal TRULI treatment, and AAV-mediated gene delivery

Doxorubicin (1 µL of 100 ng/μL) was subretinally injected into 8-week-old C57BL/6J mice to generate a doxorubicin-induced senescence model. Concurrently, 1 μL of TRULI (3.35 ng/μL) was intravitreally administered, followed by an additional intravitreal injection of 1 μL TRULI (3.35 ng/μL) three days later. The mice were euthanized one week after the initial doxorubicin injection.

An AMD-like pathology model was created by intraperitoneally injecting 25 mg/kg NaIO₃ into 8-week-old C57BL/6J mice. Retinal damage was confirmed using CFP three days after NaIO₃ administration. Subsequently, 1 μL of TRULI (16.72 ng/μL) was intravitreally injected at six-day intervals for a total of four injections. To assess the role of Yap activation via AAV-mediated expression, 8-week-old C57BL/6J mice received an intraperitoneal injection of 25 mg/kg NaIO₃, followed by a subretinal injection of 1×10⁷ TU/μL AAV2 vectors pseudotyped with an AAV8 capsid (rAAV2/8). The control mice were injected with AAV-CMV-mCherry, whereas the experimental group received AAV-CMV-mYap1[NM_001421048.1]/mCherry. The mice were euthanized two weeks post-injection to evaluate Yap activation.

TRULI (1.5 µL of 16.72 ng/μL) was intravitreally injected into 23-month-old mice to evaluate its effects in aging mice. The injections were administered four times at one-week intervals, and the mice were euthanized on day 28.

### Subretinal and intravitreal injection

For ocular injections, the eyes of mice were dilated using 5 mg/mL phenylephrine hydrochloride and 5 mg/mL tropicamide (Hanmi Pharm) and locally anesthetized with 0.5% proparacaine hydrochloride (Alcon). General anesthesia was induced via intraperitoneal injection of a mixture containing 10 mg/mL alfaxalone (Jurox), saline (Korean Pharmaceutical Industries Co., Ltd.), and xylazine (Rompun) in a 30:19:1 ratio. The anesthetized mice were positioned laterally, and their eyes were examined using an optical microscope (Olympus SZ51). A small hole was created in the sclera using a 30-gauge sterile needle (BD Bioscience). TRULI was slowly injected into the vitreous cavity for approximately 3 s using a blunt 35-gauge Hamilton microsyringe (Hamilton Company). For subretinal injections, a 30-gauge Hamilton syringe was used to create a hole in the retina through the pierced scleral site, and AAV or doxorubicin was carefully injected under the retina for approximately 3 s. The eyes were then treated with an antibiotic ophthalmic ointment (Tarivid), and the mice were kept warm until they fully recovered from anesthesia.

### Western blotting

After the mice were euthanized, their eyes were enucleated and carefully trimmed to remove surrounding muscle and fat tissue. The anterior segments and retina were removed, and RIPA lysis buffer was gently applied to the eye cup to extract proteins. The protein concentration of RPE cell lysates was measured using a BCA assay (Thermo).

Equal amounts of protein were mixed with sample buffer (Bio-Rad) and heated at 95°C for denaturation. Proteins were separated according to size using SDS-polyacrylamide gel electrophoresis (SDS-PAGE) in an acrylamide gel. Proteins were transferred onto a PVDF membrane (Millipore) at 110 volts for 80 min. The membrane was washed with Tris-buffered saline containing 0.05% Tween-20 (TBS-T) and blocked with 5% (w/v) skim milk (BD Life Sciences) in TBS-T for 1 h at RT. After blocking, the membrane was incubated overnight at 4°C with primary antibodies diluted in skim milk. After overnight incubation, the membrane was washed with TBS-T and incubated with HRP-conjugated secondary antibodies (1:5000 dilution) (Bethyl Laboratories Inc) in 5% (w/v) BSA for 90 min at RT. The membrane was washed three times for 10 min each with TBS-T. Protein bands were visualized using Pierce™ ECL Western Blotting Substrate (Thermo) and detected with the ChemiDoc image analyzer (Thermo). Each experiment was repeated at least three times to ensure reproducibility.

### Histopathologic analyses

After the mice were euthanized, their eyes, with surrounding muscle and fat tissue removed, were immediately embedded in a frozen section compound (Biosystems Inc.) and frozen at - 80°C. Frozen blocks were sectioned into 5 μm-thick slices using a cryostat (CM1860, Leica) and stained with hematoxylin and eosin (H&E). ONL thickness was quantified in the stained sections using the ImageJ 1.53 software (Fiji). Statistical analyses and plotting were performed using the GraphPad Prism 10.4.1 software (GraphPad Software).

### Immunofluorescence

The cells were fixed in 4% methanol-free formaldehyde (Thermo) for 15 min and permeabilized in 0.25% Triton X-100 (Sigma)/PBS at RT for 10 min. After washing with PBS, the samples were blocked with 10% FBS in PBS at RT for 1 h. After blocking, the tissue sections were incubated overnight at 4°C with primary antibodies diluted in a blocking buffer. The next day, the sections were washed three times with PBS. The sections were then incubated with appropriate secondary antibodies conjugated to Alexa Fluor dye (Thermo) for 1 h at RT. Nuclei were stained for 2 min with Hoechst 33342 (Thermo), followed by a final wash. Images were captured using a fluorescence microscope (Leica DMi8) and prepared using the ImageJ 1.53 software (Fiji). Statistical data were analyzed and plotted using the GraphPad Prism 10.4.1 software (GraphPad Software).

For tissue samples, OCT-embedded sections were fixed in 4% paraformaldehyde for 15 min, followed by permeabilization with 0.1% (v/v) Triton X-100 in PBS for 5 min at 4°C. After washing with PBS, the sections were blocked with 5% (v/v) BSA in PBS for 1 h at RT. The sections were then incubated overnight at 4°C with primary antibodies diluted in a blocking buffer. The next day, the sections were washed three times with PBS and incubated with Alexa Fluor dye-conjugated secondary antibodies for 1 h at RT. The nuclei were counterstained with Hoechst 33342 for 15 min at RT. After the final PBS wash, the samples were mounted using Aqua-Poly/Mount (Polysciences).

For flat-mount preparations, the cornea, lens, vitreous, and retina were carefully removed to isolate the RPE/choroid. The eyecups were fixed in 4% paraformaldehyde for 10 min and permeabilized with 0.2% (v/v) Triton X-100 in PBS for 10 min. Blocking was performed using 5% (v/v) BSA in 0.05% (w/v) Triton X-100 in PBS for 1 h at RT. Staining and mounting were performed as described above. Fluorescently stained tissues were examined under an inverted microscope (Carl Zeiss LSM 900).

### SA-β-gal assay and quantification in mouse models

Immediately after euthanizing the mice, the RPE/choroid/sclera complexes were isolated and fixed in 4% paraformaldehyde at 4°C for 1 h. The tissues were then washed with PBS and incubated overnight at 37°C in the SA-β-gal staining solution, prepared according to the manufacturer’s instructions (BioVision). After incubation, the stained RPE samples were depigmented by treatment with 7.5% H_2_O_2_ at 55°C for 40 min and then washed with PBS. The samples were mounted using Aqua-Poly/Mount (Polysciences).

Quantification of SA-β-gal staining was performed as previously described^13^. Briefly, images of SA-β-gal-stained tissues (whole mounts and cryosections) were captured and standardized to a uniform size. The ‘Split Channels’ function in ImageJ 1.53 (Fiji) was used to separate each image into red, green, and blue channels. For whole-mount samples, the red channel was selected to enhance the contrast between SA-β-gal-positive regions and the background. Stained and unstained regions were delineated using the thresholding tool, and the pixel count of SA-β-gal-positive regions was measured using the ‘Measure’ function. The percentage of the SA-β-gal-positive area was calculated by comparing the stained area to the total RPE area. For cryosectioned tissues, images were converted to the RGB color mode and inverted to improve contrast with the background. The ‘Split Channels’ function was applied, and SA-β-gal staining intensity in RPE cells was quantified. The resulting staining area and intensity data were analyzed and visualized using GraphPad Prism 5.

### Quantification of RPE cell area in mouse models

The RPE cell area was quantified using ImageJ 1.53 (Fiji). The ‘Set Scale’ function was used to calibrate pixels to μm. To emphasize cell membrane boundaries stained with ZO-1, the green channel was isolated using the ‘Split Channels’ function, and thresholding was applied to enhance contrast. The RPE cell boundaries were manually outlined using polygon selections, and the cell area (μm²) was measured with the ROI Manager. Colors were assigned to the measured values using the ‘ROI Color Coder’ function from the ‘BAR plugin’. The resulting cell area data were analyzed and visualized using GraphPad Prism 5.

### Color fundus photography and fundus autofluorescence imaging

Color fundus photographs (CFP) of the mouse eyes were captured after pupil dilation using a TRC-50 IX camera (Topcon) connected to a Nikon digital imaging system. Fundus autofluorescence imaging was performed using an HRA + OCT Spectralis system (Heidelberg Engineering).

### Electroretinography (ERG) Analysis

The mice were adapted to the dark overnight. Before the ERG measurements, 5 mg/mL phenylephrine hydrochloride and 5 mg/mL tropicamide (Hanmi Pharm) were administered to dilate the pupils. Anesthesia was induced by an intraperitoneal injection of 10 mg/mL alfaxalone (Jurox) and xylazine (Rompun). The mice were placed on a 37°C heating pad to maintain their body temperature during the procedure, and 2% hypromellose (Samil) was applied to prevent corneal dehydration. A corneal electrode was carefully positioned on the eyes for ERG recording using the Diagnosys Celeris System (Diagnosys). Scotopic full-field ERG (Ganzfeld ffERG) responses, including a- and b-waves, were recorded using white flashes at intensities from 0.001 to 10 cd·s/m², with three sweeps averaged per intensity. Standard ERG responses were obtained for both eyes by recording three sweeps from two channels. The c-wave was recorded using a stimulus intensity of 150.0 cd·s/m², and its amplitude was measured from the maximal negativity of the c-wave to the peak of the c-wave.

### Single-cell RNA sequencing analysis

For scRNA-seq, young and old mice were euthanized to dissociate the cells from the RPE. Their eyes were immediately enucleated and the anterior eye cups were dissected. After carefully removing the retina, the RPE cells were dissociated according to the manufacturer’s instructions for the papain dissociation system (Worthington). Briefly, RPE cells were carefully dissociated using a papain solution (20 U/ml) supplemented with DNase-I (2000 U/ml) in Earle’s balanced salt solution (EBSS) and incubated in a water bath at 37 °C for 30 min. The dissociated single RPE cell pellet was washed with PBS and collected by centrifugation at 1200 rpm for 3 min. The RPE samples were then filtered through a 30 µm cell strainer (Miltenyi Biotech) and washed twice with cold Ca²⁺- and Mg²⁺-free 0.04% BSA/PBS at 300 *g* for 5 min at 4°C. The samples were gently resuspended in cold Ca²⁺- and Mg²⁺-free 0.04% BSA/PBS and counted using a LUNA-FX7™ Automated Fluorescence Cell Counter (Logos Biosystems) after acridine orange (AO) and propidium iodide (PI) staining (Logos Biosystems).

scRNA-seq libraries were prepared using the 10x Chromium Controller and Next GEM Single Cell 5’ Reagent v2 kits (10x Genomics), following the 10x Chromium Single Cell 5’ v2 protocol. Cell suspension with a target recovery of 10,000 cells was combined with a reverse transcription master mix and loaded onto a Single Cell K-Chip (10x Genomics) with single cell 5′ gel beads and partitioning oil to generate gel beads-in-emulsion (GEMs), in which RNA transcripts from individual cells were uniquely barcoded and reverse transcribed. The resulting barcoded full-length cDNA was amplified by PCR, and for 5’ Gene Expression Library preparation, the cDNA underwent enzymatic fragmentation, end repair, A-tailing, adapter ligation, and index PCR. The purified libraries were quantified using the qPCR Qualification Protocol Guide (KAPA), evaluated for quality using an Agilent 4200 TapeStation (Agilent Technologies), and sequenced on an Illumina HiSeq platform (Illumina).

Gene expression matrices were generated using Cell Ranger 6.0.1 (10x Genomics) with the mouse genome reference data (mm10). Quality control filtering was performed using cells with an nFeature_RNA > 200 and less than 10% mitochondrial content. Doublet cells were removed using Scrublet (version 0.2.3) with a doublet score threshold of < 0.1; scrub_doublets(min_counts = 2, min_cells = 3, min_gene_variability_pctl = 85, n_prin_comps = 30). After filtering, 32,285 gene expression matrices were obtained from the 27,840 cells.

To cluster the old and young mice samples together, we used the Seurat package (version 4.1.1). Clustering and visualization were performed using Uniform Manifold Approximation and Projection (UMAP), based on the top 25 principal components (PCs) and a clustering resolution of 0.4. To subset the data into RPE subclusters (clusters 2, 3, 6, 12, 14, and 15), subsetting was performed by comparing the expression of known RPE marker genes, with the top 11 PCs and a clustering resolution of 0.4.

To identify the dominance of old or young cells in each cluster, dominance analysis was conducted by calculating the old or young cell ratio as

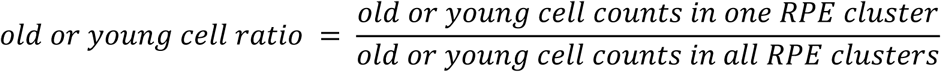

throughout all RPE clusters, with a ratio of two or more being identified as either old- or young-dominant. Differential expression analysis between the old- and young-dominant groups was performed using Seurat’s FindMarkers function, with significance defined as |log_2_FC| ≥ 0.5 and adjP < 0.05. The enriched pathways of differential expression gene (DEG) sets were identified using the GO cellular component and KEGG databases using enrichR (version 3.1). To identify the relationship between samples and a specific gene set, the gene signature score was calculated using AddModuleScore from the Seurat package; AddModuleScore (object, list(features), assay = “RNA”). The Wilcoxon test was used to compare gene expressions in violin plots. All packages used in the analysis were implemented using the R programming software (version 4.1.1).

For human scRNA-seq analysis, raw fastq files (Table S3) were downloaded and processed to generate gene expression matrices using Cell Ranger 6.1.2 (10x Genomics) with the human genome reference data (hg38 and gencode v46). Single cell quality control filtering was performed using the following criteria: nFeature_RNA > 200, nFeature_RNA < 6000, nCount_RNA > 200, log10 (nFeature_RNA/nCount_RNA) > 0.8, and mitochondrial content < 30%. After filtering, 63,086 gene expression matrices were obtained from the 83,740 cells.

To correct batch effect and integrate the data sets for visualization, we performed harmony integration using the ‘IntegrateLayers’ function in the Seurat package (version 5.2.0). To annotate cells, we first transferred cell labels from the tabular Sapiens eye scRNA-seq data^33^ using Seurat’s FindTransferAnchors and TransferData functions. We then annotated clusters using the cell labels comprising more than 70% of the cells within the cluster. If there were no transferred cell labels representing more than 70% of the Seurat clusters, we manually checked the expression of cell type marker genes to annotate the cell types (mast cells, macrophages, Schwann cells, and photoreceptors). Finally, 4,721 RPE cells were isolated for differential expression analysis. Differential expression analysis (AMD vs. normal adult and adult vs. fetal) was performed using Seurat’s FindMarkers function, with significance defined as |log2FC| ≥ 1 and adjP < 0.05. Enriched pathways of differential expression gene (DEG) sets were identified from the GO cellular component and KEGG databases using ShinyGO (version 0.82).

### Bulk RNA sequencing analysis

ARPE19 cells expressing control vectors were cultured on plastic or 1.5 kPa, and ARPE19 cells overexpressing YAP-HA were cultured on 1.5 kPa as previously described. Cells were harvested after 5 days and the total RNA was isolated using QIAzol Lysis Reagent (QIAGEN). Total RNA concentration was measured using the Quant-IT RiboGreen Assay (Invitrogen). Samples were analyzed using the TapeStation RNA ScreenTape system (Agilent) to assess RNA integrity, and only RNA samples with an RNA Integrity Number (RIN) greater than 7.0 were selected for RNA library preparation. For each sample, 1 µg of total RNA was used to construct libraries using the Illumina TruSeq Stranded mRNA Sample Prep Kit (Illumina). Poly-A mRNA was isolated using poly-T-coated magnetic beads and fragmented at high temperatures in the presence of divalent cations. A cDNA library was generated from fragmented RNA using SuperScript II reverse transcriptase (Invitrogen). The prepared libraries were quantified using the KAPA Library Quantification Kit for Illumina platforms according to the manufacturer’s protocol (KAPA BIOSYSTEMS). Library quality was assessed using TapeStation D1000 ScreenTape (Agilent). The indexed libraries were then sequenced on an Illumina NovaSeqX platform (Illumina) with paired-end 2×100 bp reads performed by Macrogen Inc.

All raw files were mapped to human genome reference data (GRCh38) using STAR (version 2.7.3a). After evaluating the quality of the raw data using FastQ Screen (version 0.14.1), we calculated TPM (transcripts per million) using RSEM (RNA-Seq by Expectation-Maximization) (version 1.3.1). To determine the differences among the three samples, pairwise DEG analysis was conducted using DESeq2 (version 1.34.0) with genes having an average TPM value of 5 or more. In addition, we calculated the Z-score of the upregulated or downregulated DEGs and then compared the DEG patterns of the three samples in a heatmap. All plots for visualization were generated using ggplot2 in the R package (version 4.1.1).

For mouse samples, aged mice treated with or without TRULI were euthanized for RNA extraction, and their eyes were immediately enucleated. The eyeballs were trimmed to remove the surrounding muscles, and the anterior segments were dissected. After carefully isolating the retina, lysis buffer was gently applied to the eye cup, and RNA was extracted using the RNeasy Mini Kit (QIAGEN) according to the manufacturer’s instructions. The sequencing library was constructed using the SMARTer Stranded RNA-Seq Kit according to the manufacturer’s instructions. Library quantification, quality assessment, and sequencing were performed on the Illumina NovaSeqX platform as described above. STAR (version 2.7.10b) was used to map raw sequencing data to the mouse genome reference data (GRCm39). FastQ Screen (version 0.14.1) was used to check the quality of the raw data.

TPM was obtained using RSEM (version 1.3.1). DEG analysis was performed with NOISeqbio (version 2.38.0) using genes with an average TPM value of 5 or more.

**Fig. S1.**
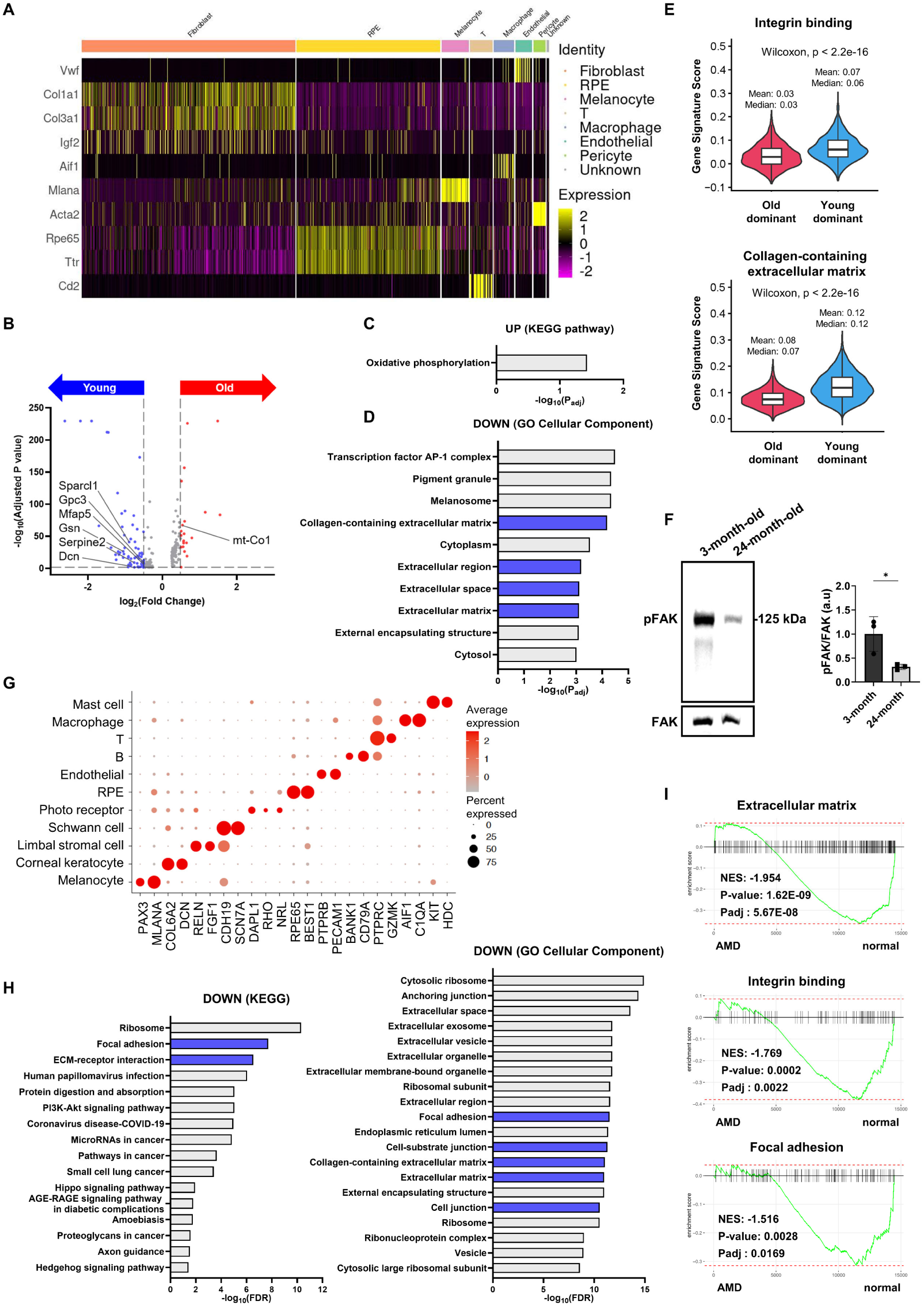
scRNA-seq analysis of aged mouse and AMD human RPE cells uncovers impaired cell-matrix adhesion. (A) Identification of cell types using marker gene expression. (B) Volcano plot of DEGs in RPE cells from 3-month-old and 24-month-old mice. The red and blue dots indicate either upregulated or downregulated genes in old mouse RPE, with cutoff values for DEGs: |log_2_FC| ≥ 0.5, adjP < 0.05. (C and D) The top pathways ranked by adjusted p-values from GO analysis with upregulated (C) and downregulated (D) genes from (B). (E) Violin plots showing gene scores of old-dominant and young-dominant clusters from two different gene sets: integrin binding and collagen-containing extracellular matrix. (F) Western blot analysis of phosphorylated and total FAK proteins in RPE cells from 3-month and 24-month old mice. *n* = 3 samples per group. (G) Dot plots showing the expression of tissue-specific marker genes in human eye scRNA-seq clusters. The red color indicates high expression, and the dot size indicates the percentage of cells showing gene expression. (H) The top pathways ranked by FDR from GO analysis with downregulated genes in adult RPE compared to fetal RPE. (I) Gene set enrichment analysis (GSEA) with human RPE scRNA-seq data. Genes were ranked by expression fold change in AMD RPE compared to normal RPE. Two-sided Student’s t-test, *P < 0.05. Exact *P* values are listed in Table S10.

**Fig. S2.**
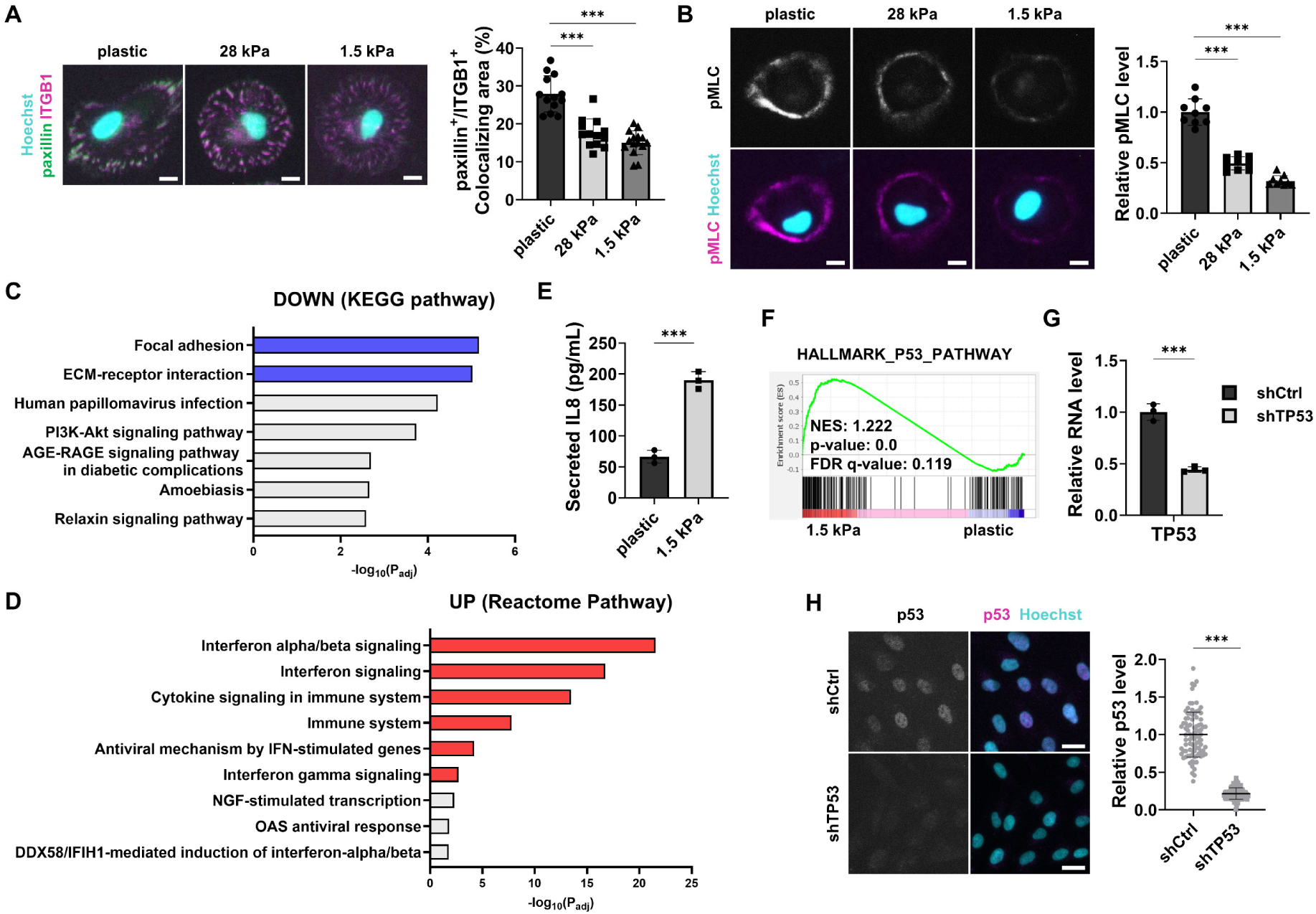
ARPE19 cells cultured on soft substrate show senescence phenotypes. (A) Immunofluorescence assay for paxillin and ITGB1 in ARPE19 cells cultured on substrate with different stiffness. *n* = 13 cells for Plastic, *n* = 13 cells for 28 kPa, *n* = 15 cells for 1.5 kPa from three independent experiments. The percentages of colocalizing area were analyzed by paxillin and ITGB1 double positive area over total ITGB1 area. (B) Immunofluorescence assay for pMLC in ARPE19 cells cultured on substrate with different stiffness. *n* = 9 regions from three independent experiments. (C and D) The top pathways ranked by adjusted p-values from GO analysis with downregulated (C) and upregulated (D) genes in ARPE19 cultured on 1.5 kPa substrate. (E) Enzyme-linked immunosorbent assay (ELISA) for IL8 in the media of ARPE19 cells cultured either on plastic or 1.5 kPa substrate. *n* = 3 from three independent experiments. (F) GSEA analysis of HALLMARK_P53_PATHWAY from MSigDB with the RNA-seq data from ARPE19 cultured either on plastic or 1.5 kPa substrate. (G and H) Validation of lentiviral vector expressing TP53 shRNA in ARPE19 cells by qPCR (G) and immunostaining (H). *n* = 3 samples for qPCR (G) and *n* = 90 cells for immunostaining (H) from three independent experiments. Two-sided Student’s t-test, ***P < 0.001. Exact *P* values are listed in Table S10. Scale bars: 10 μm (A, B), 25 μm (H)

**Fig. S3.**
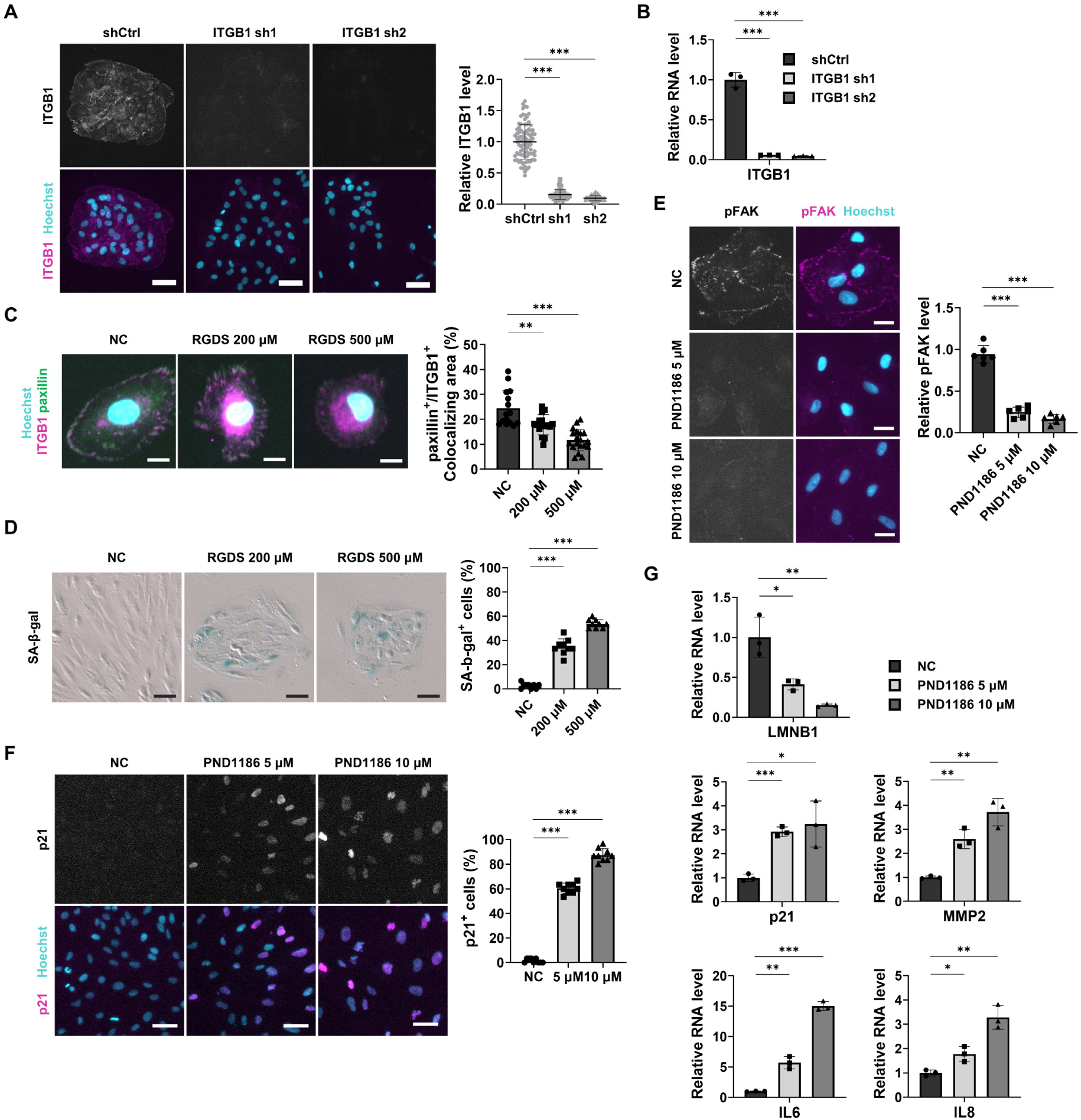
Integrin signaling inhibition induces RPE senescence. (A and B) Validation of lentiviral vectors expressing ITGB1 shRNAs in ARPE19 cells by immunostaining (A) and qPCR analysis (B). *n* = 90 cells for immunostaining, *n* = 3 samples for qPCR from three independent experiments. (C) Immunofluorescence assay for paxillin and ITGB1 in ARPE19 cells plated and cultured with RGDS for 96 hours. *n* = 15 cells for NC, *n* = 16 cells for RGDS 200 μM, and *n* = 19 cells for RGDS 500 μM from three independent experiments. The percentages of colocalizing area were analyzed by paxillin and ITGB1 double positive area over total ITGB1 area. (D) SA-β-gal staining in ARPE19 cells plated and cultured with RGDS for 96 hours. *n* = 9 regions from three independent experiments. (E and F) Immunofluorescence assay for pFAK (E), and p21 (F) in ARPE19 cells treated with PND1186 either for 24 hours (E) or 96 hours (F). *n* = 6 regions for pFAK (E) from two independent experiments, *n* = 9 regions for p21 (F) from three independent experiments. (G) qPCR analysis of LMNB1, p21, and SASP genes in ARPE19 cells treated with PND1186 for 96 hours. *n* = 3 samples from three independent experiments. Two-sided Student’s t-test, ***P < 0.001, **P < 0.01, *P < 0.05. Exact *P* values are listed in Table S10. Scale bars: 10 μm (C), 25 μm (E), 50 μm (A, G)

**Fig. S4.**
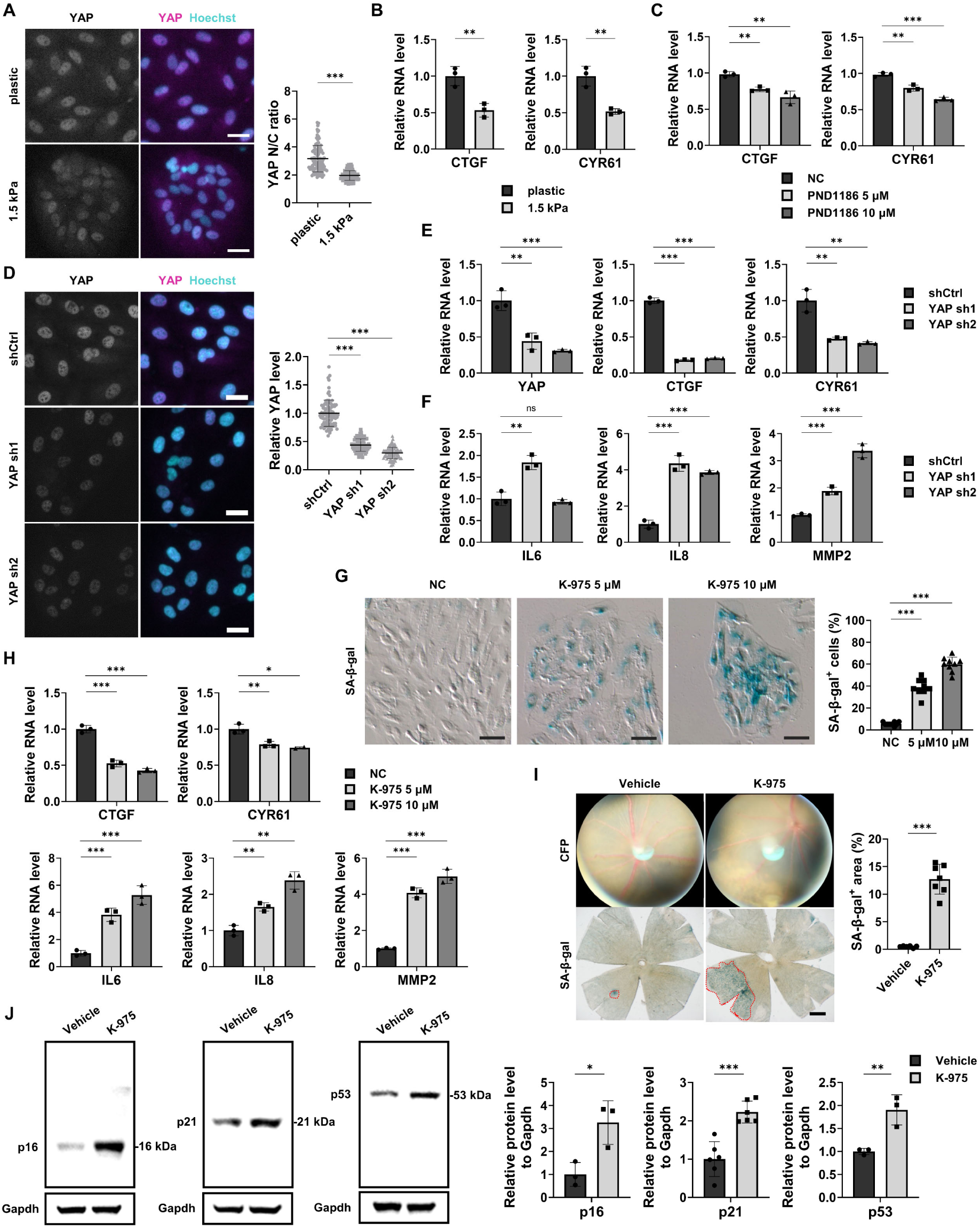
RPE senescence is regulated by YAP. (A and B) Immunofluorescence assay for YAP (A) and qPCR analysis for YAP target genes (B) in ARPE19 cells cultured on either plastic or 1.5 kPa. YAP activity was quantified by the ratio of nuclear over cytoplasmic intensities (N/C ratio). *n* = 90 cells for immunostaining (A) and *n* = 3 samples for qPCR (B) from three independent experiments. (C) qPCR analysis for YAP target genes in ARPE19 cells treated with PND1186 for 96 hours. *n* = 3 samples from three independent experiments. (D and E) Validation of lentiviral vectors expressing YAP shRNAs in ARPE19 cells by immunostaining (D) and qPCR analysis (E). *n* = 90 cells for immunostaining (D), *n* = 3 samples for qPCR (E) from three independent experiments. (F) qPCR analysis of SASP genes in ARPE19 cells after YAP KD. *n* = 3 samples from three independent experiments. (G) SA-β-gal staining in ARPE19 cells treated with K-975 for 120 hours. *n* = 9 regions from three independent experiments. (H) qPCR analysis of YAP target genes and SASP genes in ARPE19 cells treated with K-975 for 120 hours. *n* = 3 samples from three independent experiments. (I) Representative color fundus photographs (CFPs) and SA-β-gal staining of RPE/choroid flat mounts of C57BL/6J mice subretinally injected with K-975 for 1 week. *n* = 6 mice for Vehicle and *n* = 7 mice for K-975. (J) Western blot analysis of isolated RPE cells from mouse eyes for p16, p21, and p53 in K-975-treated mice. *n* = 3 samples per group for p16 and p53; *n* = 6 samples for p21. Two-sided Student’s t-test, ***P < 0.001, **P < 0.01, *P < 0.05, ns; not significant. Exact *P* values are listed in Table S10. Scale bars: 25 μm (A, D), 50 μm (G), 500 μm (I)

**Fig. S5.**
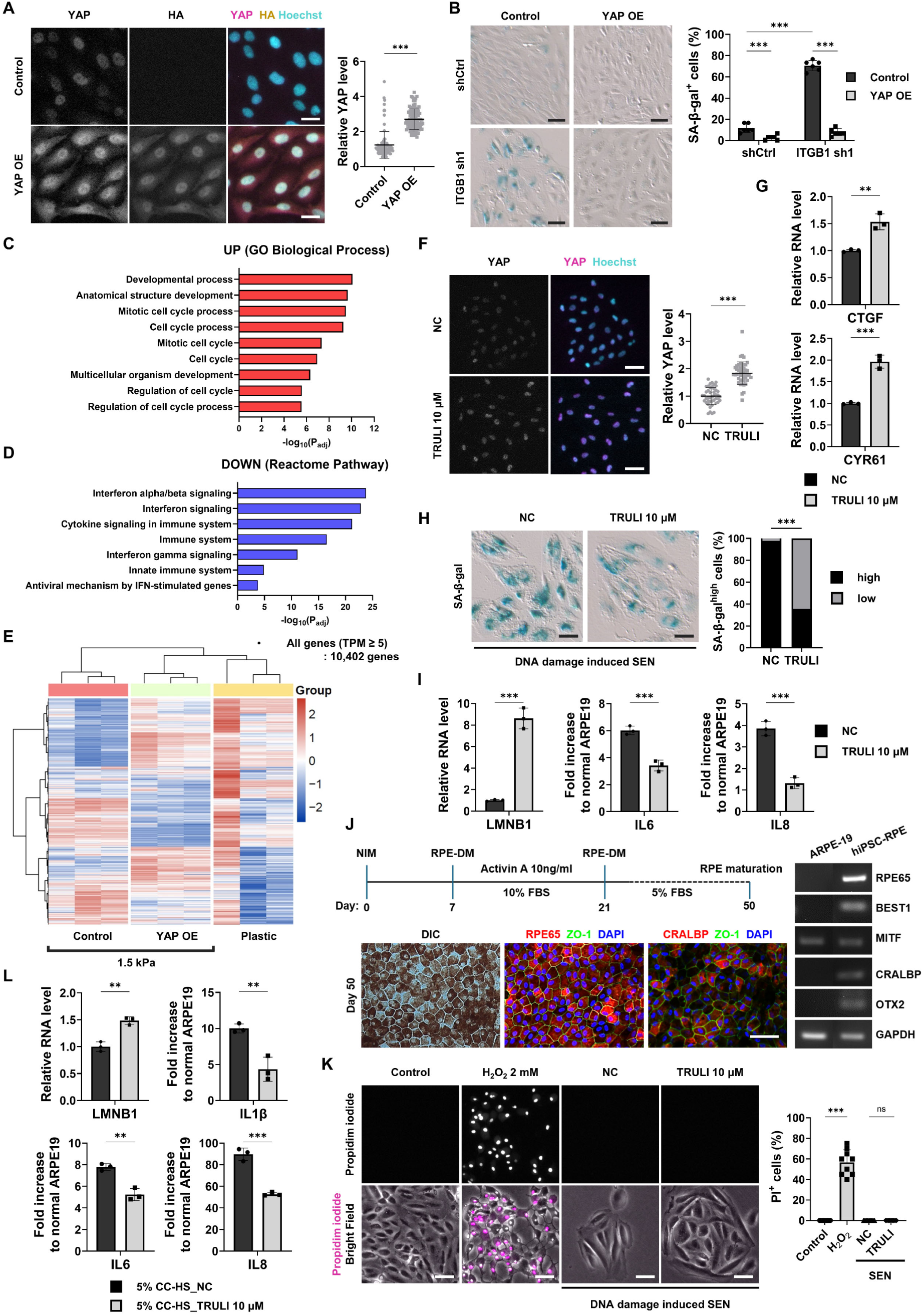
YAP activation reverses senescence phenotypes *in vitro*. (A) Immunostaining for YAP and HA tag in ARPE19 cells after transduction with lentiviral vectors overexpressing YAP-HA. *n* = 90 cells from three independent experiments. (B) SA-β-gal staining in YAP overexpressing ARPE19 cells followed by ITGB1 KD. *n* = 6 regions from two independent experiments. (C and D) The top pathways ranked by adjusted p-values from GO analysis with upregulated (C) and downregulated (D) genes in YAP overexpressing ARPE19 cells cultured on 1.5 kPa substrate. e. Hierarchical clustering analysis of ARPE19 cells cultured on plastic or 1.5 kPa substrate after transduction with lentiviral vectors overexpressing YAP. (F and G) Immunostaining for YAP (F) and qPCR analysis of YAP target genes (G) in ARPE19 cells treated with TRULI for 24 hours. *n* = 45 cells for immunostaining (F) from two independent experiments, *n* = 3 samples for qPCR (G) from three independent experiments. (H and I) SA-β-gal staining (H), and qPCR analysis of LMNB1 and SASP genes (I) after TRULI treatment for 24 hours in senescent ARPE19 cells induced by Doxorubicin treatment. *n* = 9 regions for SA-β-gal staining (H) and *n* = 3 samples for qPCR (I) from three independent experiments. (J) Schematic representation of the differentiation process from hiPSCs to RPE. RPE maturation is validated by the morphology and RPE-specific marker expression at 50 days post-differentiation. (K) Propidium iodide (PI) staining of ARPE19 cells treated with 2 mM H_2_O_2_ for 12 hours or Doxorubicin-induced senescent ARPE19 treated with TRULI for 24 hours. *n* = 9 regions from three independent experiments. (L) qPCR analysis of LMNB1 and SASP genes in CC-HS induced senescent ARPE19 cells treated with TRULI for 24 hours. *n* = 3 samples from three independent experiments. Two-sided Student’s t-test and two-way ANOVA, ***P < 0.001, **P < 0.01, *P < 0.05, ns; not significant. Exact *P* values are listed in Table S10. Scale bars: 25 μm (A), 50 μm (B, G, H, K)

**Fig. S6.**
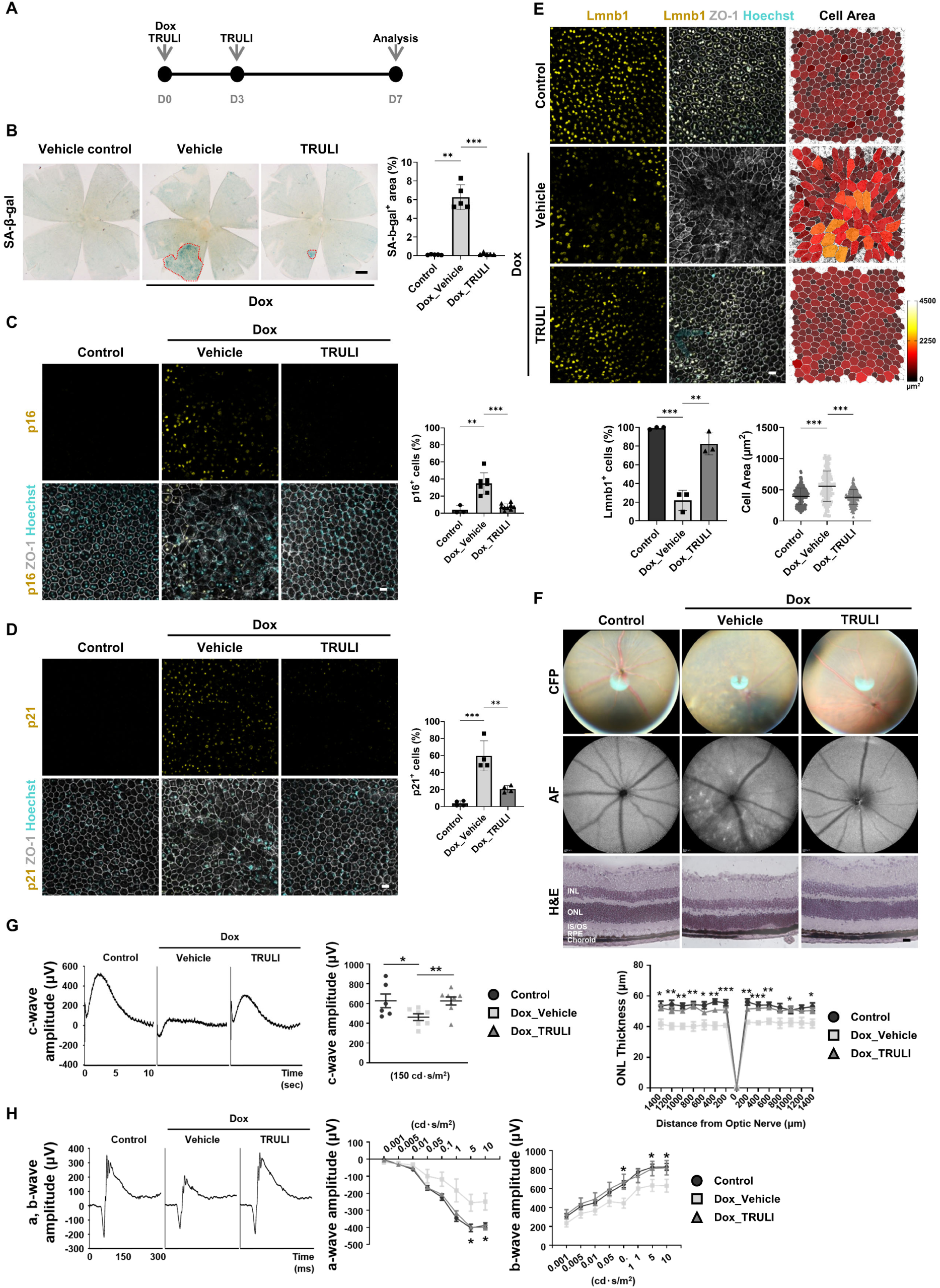
Yap activation by TRULI recovers visual function in a Doxorubicin-induced senescence mouse model. (A) Experimental scheme for evaluating TRULI effects on visual function in a Doxorubicin-induced senescence mouse model. (B) SA-β-gal staining in RPE/choroid flat mounts. *n* = 5 mice for control and Dox_Vehicle, *n* = 6 mice for Dox_TRULI. (C and D) Immunostaining for p16 (C), p21 (D), and ZO-1 in RPE/choroid flat mounts. *n* = 3 mice for control, *n* = 7 mice for Dox_Vehicle, *n* = 9 mice for Dox_TRULI in (C); *n* = 4 mice per group in (D). (E) Immunostaining for Lmnb1 in RPE/choroid flat mounts. *n* = 3 mice per group. The measured cell areas were visualized using color mapping with the LUT Red Hot scale. *n* = 224 cells for control, *n* = 154 cells for Dox_Vehicle, n = 227 cells for Dox_TRULI. (F) CFP, autofluorescence (AF) images, and H&E-stained retinas. The thickness of the outer nuclear layer (ONL) was measured at 200 μm intervals from the center of the optic nerve. *n* = 5 mice per group. (G and H) ERG analysis in Doxorubicin-injected mice treated with TRULI. The amplitude of the c-wave (μV) was measured at 150 cd·s/m² (G). *n* = 6 mice for control, *n* = 7 mice for Dox_Vehicle, and *n* = 8 mice for Dox_TRULI. The amplitude of the scotopic a-wave and b-wave was measured across the range of 0.001-10 cd·s/m² (H). *n* = 4 mice per group (a-wave); *n* = 6 mice for control, *n* = 4 mice for Dox_Vehicle and Dox_TRULI (b-wave). Two-sided Student’s t-test, ***P < 0.001, **P < 0.01, *P < 0.05. Exact *P* values are listed in Table S10. Abbreviations: INL, inner nuclear layer; ONL, outer nuclear layer; IS, inner segment of photoreceptors; OS, outer segment of photoreceptors. Scale bars: 20 μm (C, D, E, F), 500 μm (B)

**Fig. S7.**
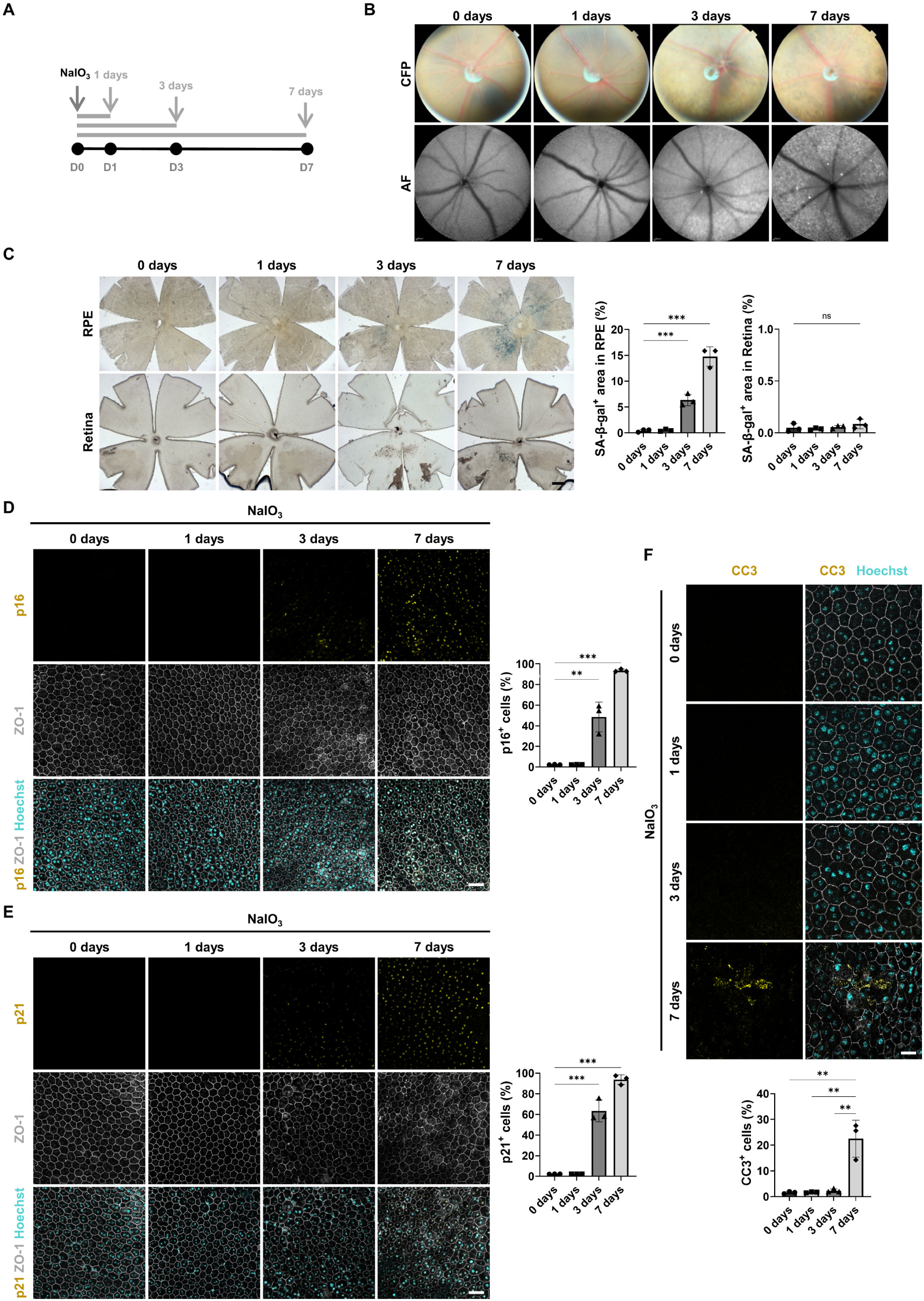
NaIO_3_ induces senescence in RPE cells *in vivo*. (A) Experimental scheme for evaluating RPE senescence in NaIO_3_-treated mice. (B and C) CFPs and AF images (B) and SA-β-gal staining images (C) in RPE/choroid and retina flat mounts. *n* = 3 samples per group. (D and E) Immunostaining for p16 (D), p21 (E), and ZO-1 in RPE/choroid flat mounts. *n* = 3 samples per group. (F) Immunostaining for cleaved caspase-3 (CC3) in RPE/choroid flat mounts. *n* = 3 samples per group. Two-sided Student’s t-test: ***P < 0.001, **P < 0.01, ns; not significant. Exact *P* values are listed in Table S10. Scale bars: 10 μm (F), 20 μm (D, E), 500 μm (C).

**Fig. S8.**
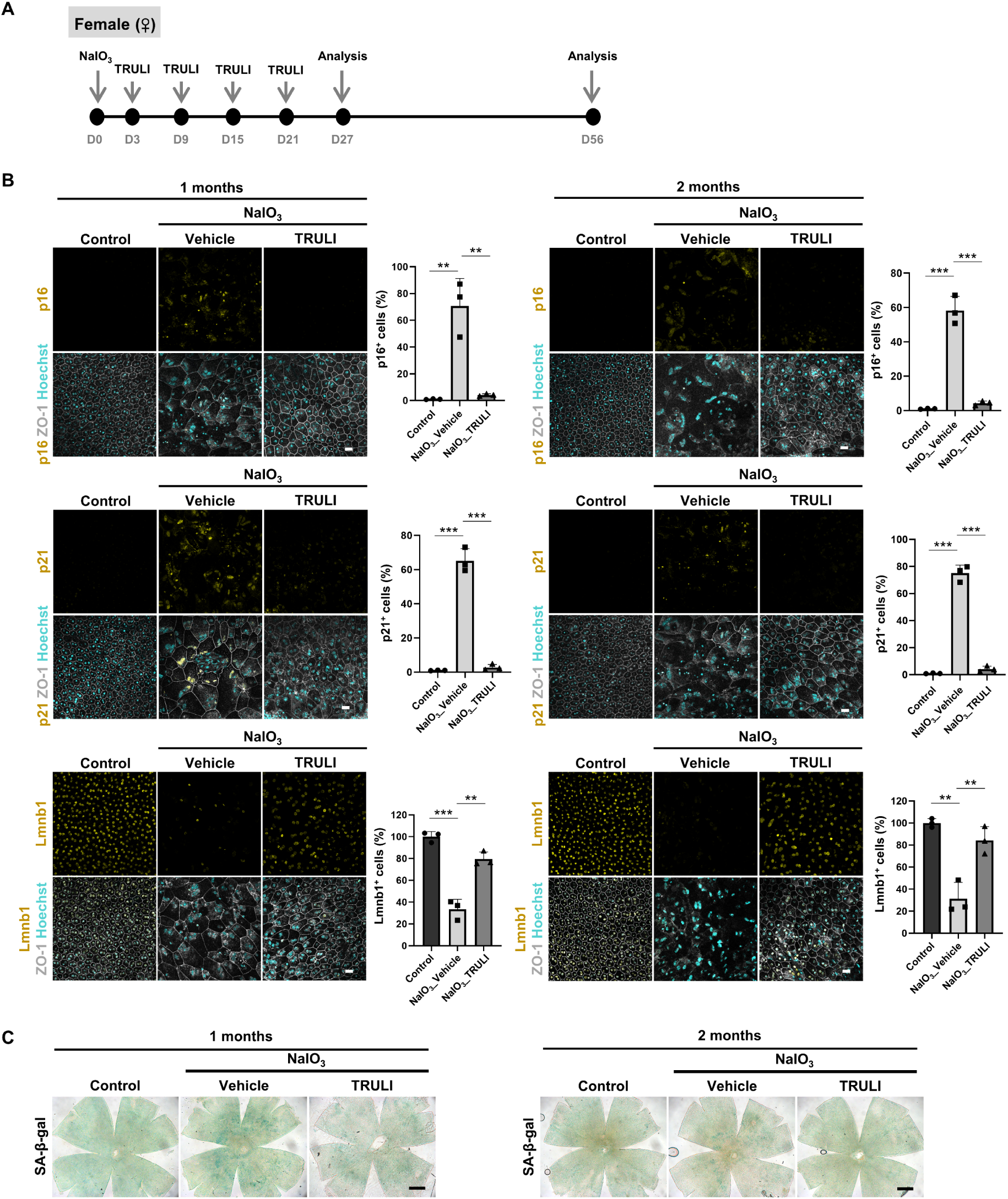
Intravitreal TRULI treatment alleviates RPE senescence and preserves tissue morphology in female NaIO3-induced AMD-like mice. (A) Experimental timeline of NaIO₃ and intravitreal TRULI administration in female C57BL/6J mice, with analysis at 1 and 2 months post-injection. (B) Immunofluorescence images and quantification of RPE/choroid flat mounts stained for p16, p21, and Lmnb1 at 1 month (left) and 2 months (right) after NaIO₃ injection. n = 3 mice per group. (C) SA-β-gal staining images of RPE/choroid flat mounts. Two-sided Student’s t-test, ***P < 0.001, **P < 0.01. Exact P values are listed in Table S10. Scale bars: 20 μm (B), 500 μm (C).

**Fig. S9.**
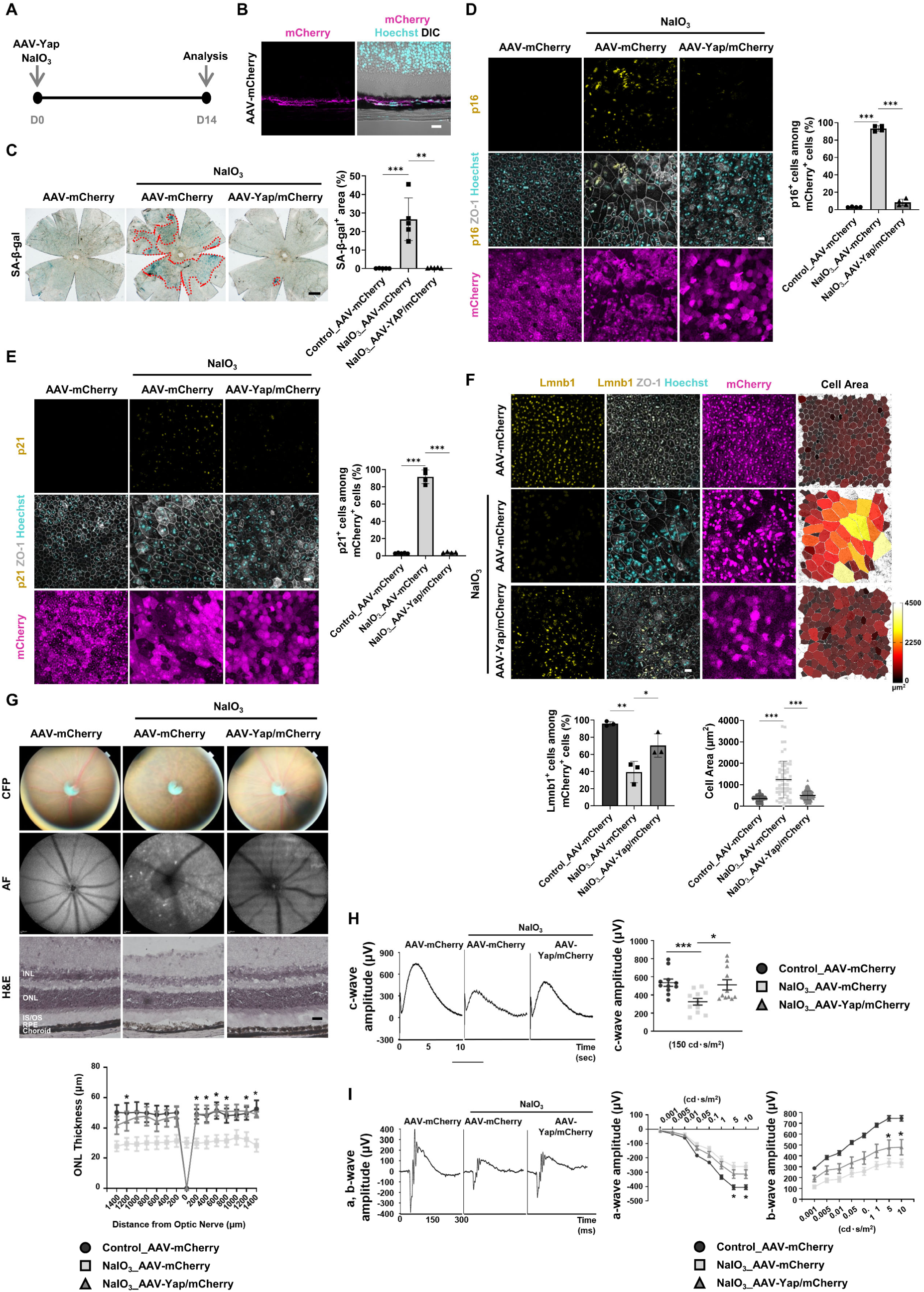
Yap overexpression through AAV-mediated gene delivery restores visual function *in vivo*. (A) Experimental scheme for evaluating the effects of AAV-Yap on visual function in an AMD-like pathology mouse model. (B) Expression patterns of mCherry in the RPE and retina two weeks after subretinal injection of AAV-mCherry (1x10⁷ TU/μL). (C) SA-β-gal staining in RPE/choroid flat mounts after AAV-Yap injection. *n* = 5 samples per group. (D, E, and F) Immunostaining for p16 (D), p21 (E), Lmnb1 (F), and ZO-1 in RPE/choroid flat mounts. *n* = 4 mice per group (D); *n* = 5 mice for control_AAV-mCherry, *n* = 4 mice for NaIO_3__AAV-mCherry and NaIO_3__AAV-Yap/mCherry in (E); *n* = 3 mice per group in (F). The measured cell areas (F) were visualized using color mapping with the LUT Red Hot scale. *n* = 242 cells for control_AAV-mCherry, *n* = 61 for NaIO_3__AAV-mCherry, *n* = 173 for NaIO_3__AAV-Yap/mCherry. (G) CFP, AF images, and H&E-stained retinas. ONL thickness was measured at 200 μm intervals from the optic nerve center. *n* = 3 mice for control_AAV-mCherry, *n* = 4 mice for NaIO_3__AAV-mCherry, and *n* = 3 mice for NaIO_3__AAV-Yap/mCherry. (H and I) ERG analysis in NaIO_3_-induced AMD mice injected with AAV. The amplitude of the c-wave (μV) was measured at 150 cd·s/m² (H). *n* = 11 mice per group. The amplitude of the scotopic a-wave and b-wave was measured across the range of 0.001-10 cd·s/m² (I). *n* = 12 mice for control_AAV-mCherry and NaIO_3__AAV-mCherry, *n* = 13 mice for NaIO_3__AAV-Yap/mCherry (a-wave); *n* = 9 mice for control_AAV-mCherry, *n* = 7 mice for NaIO3_AAV-mCherry and NaIO_3__AAV-Yap/mCherry (b-wave). Two-sided Student’s t-test, ***P < 0.001, **P < 0.01, *P < 0.05. Exact *P* values are listed in Table S10. Scale bars: 10 μm (B), 20 μm (D, E, F, G), 500 μm (C)

**Figure S10.**
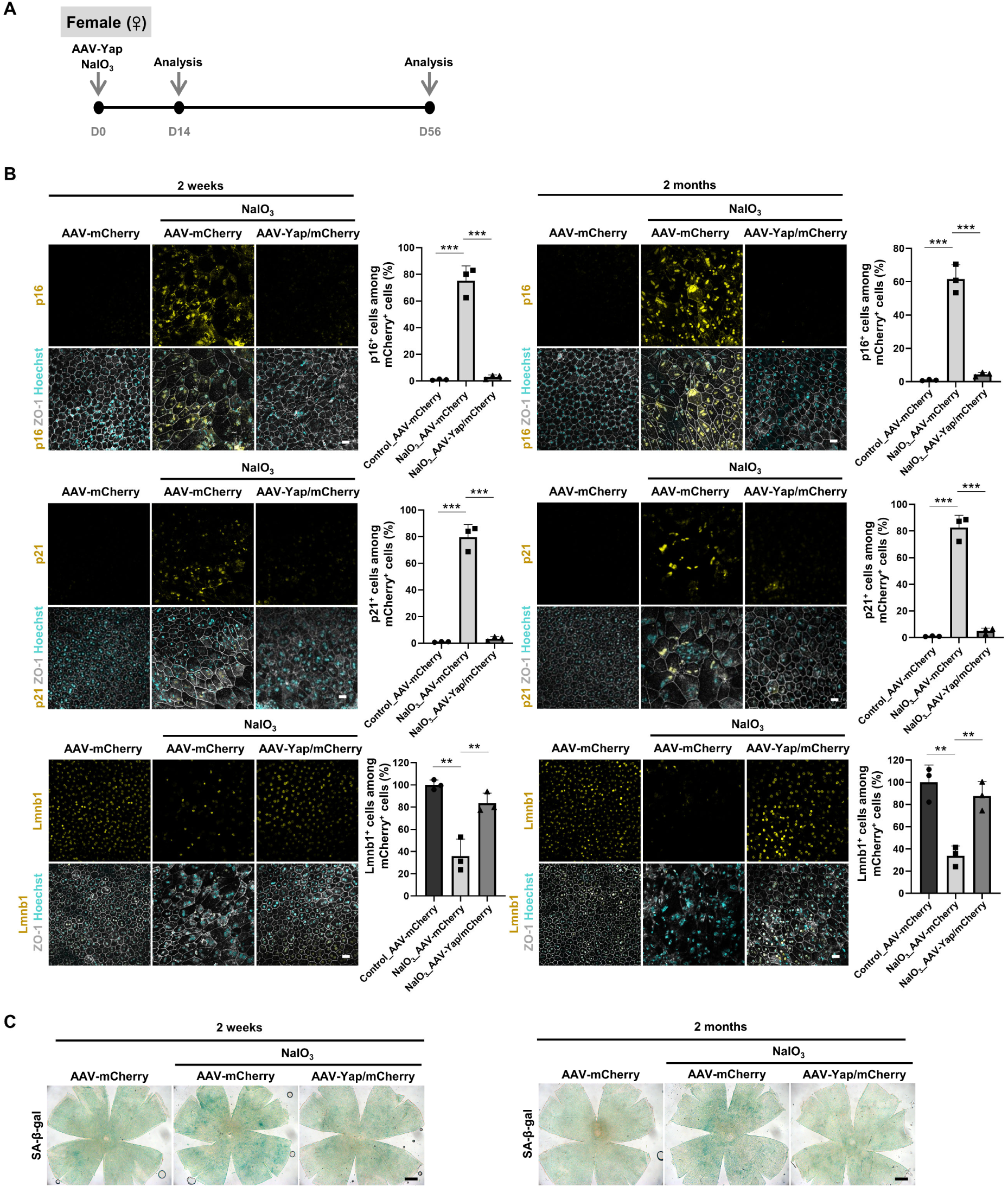
Subretinal AAV-Yap delivery reduces RPE senescence and preserves tissue integrity in female NaIO₃-induced AMD-like mice. (A) Experimental timeline of NaIO₃ injection followed by subretinal delivery of AAV-Yap or control AAV-mCherry in female C57BL/6J mice, with analyses at 2 weeks and 2 months post-injection. (B) Immunofluorescence images and quantification of RPE/choroid flat mounts stained for p16, p21, and Lmnb1 in AAV-Yap-treated mice at both 2-week (left) and 2-month (right) timepoints. n = 3 mice per group. (C) SA-β-gal staining images of RPE/choroid flat mounts. Two-sided Student’s t-test, ***P < 0.001, **P < 0.01. Exact P values are listed in Table S10. Scale bars: 20 μm (B), 500 μm (C).

## SUPPLEMENTAL TABLE LEGENDS

**Table S1. DEGs in RPE cells from young and old mice eyes, related to Figure S1**

**Table S2. DEGs in RPE cells from young dominant and old dominant clusters, related to Figure 1**

**Table S3. Published scRNA-seq datasets used in this study**

**Table S4. DEGs in RPE cells from fetal and adult human eyes, related to Figure S1**

**Table S5. DEGs in RPE cells from normal adult and AMD patient human eyes, related to Figure 1**

**Table S6. DEGs in ARPE19 cells cultured either on plastic or 1.5 kPa hydrogel substrate, related to Figure 2**

**Table S7. DEGs in ARPE19 cells cultured on 1.5 kPa hydrogel substrates after YAP overexpression, related to Figure 5**

**Table S8. DEGs in RPE cells from old mice treated with TRULI, related to Figure 7**

**Table S9. qPCR Primers used in this study**

**Table S10. *P* values reported in this study**

